# Methylomic profiles reveal sex-specific shifts in leukocyte composition associated with post-traumatic stress disorder

**DOI:** 10.1101/594127

**Authors:** Grace S. Kim, Alicia Smith, Fei Xu, Vasiliki Michopoulos, Adriana Lori, Don L. Armstrong, Allison E. Aiello, Karestan C. Koenen, Sandro Galea, Derek E. Wildman, Monica Uddin

## Abstract

Post-traumatic stress disorder (PTSD) is a debilitating mental disorder precipitated by trauma exposure. However, only some persons exposed to trauma develop PTSD. There are sex differences in risk; twice as many women as men, develop a lifetime diagnosis of PTSD. Methylomic profiles derived from peripheral blood are well-suited for investigating PTSD because DNA methylation (DNAm) encodes individual response to trauma and may play a key role in the immune dysregulation characteristic of PTSD pathophysiology. In the current study, we leveraged recent methodological advances to investigate sex-specific shifts in DNAm-based leukocyte composition that are associated with lifetime PTSD. We estimated leukocyte composition on a combined methylation array dataset (483 participants, ∼450k CpG sites) consisting of two civilian cohorts, the Detroit Neighborhood Health Study and Grady Trauma Project. Sex-stratified Mann-Whitney U test and two-way ANCOVA revealed that lifetime PTSD was associated with a small but significant elevation in monocyte proportions in males, but not in females (Holm-adjusted p-val < 0.05). No difference in monocyte proportions was observed between current and remitted PTSD cases in males, suggesting that this sex-specific peak shift reflects a long-standing trait of lifetime history of PTSD, rather than current state of PTSD. Associations with lifetime PTSD or PTSD status were not observed in any other leukocyte subtype and our finding in monocytes was confirmed using cell estimates based on a different deconvolution algorithm, suggesting that our sex-specific findings are robust across cell estimation approaches. Overall, our main finding of elevated monocyte proportions in males, but not in females with lifetime history of PTSD provides evidence for a sex-specific shift in peripheral blood leukocyte composition that is detectable in methylomic profiles and that reflects long-standing changes associated with PTSD diagnosis.

## 1. INTRODUCTION

Post-traumatic stress disorder (PTSD) is a debilitating mental disorder that is precipitated by a traumatic experience involving direct or indirect exposure to actual or threatened death, serious injury, or sexual violence^1^. PTSD presents with intrusive and persistent re-experiencing of the traumatic event, avoidance of distressing, trauma-associated stimuli, negative alterations in cognition and mood, and alterations in arousal/reactivity that persist for longer than a month^1^. While most individuals are exposed to a potentially traumatic event at some point in their lives, only some develop PTSD^2–8^, suggesting that the disorder reflects a distinct inability to reinstate homeostasis after psychological trauma in vulnerable individuals^9^.

Epidemiological studies have identified sex to be a significant vulnerability factor for developing PTSD, with women twice as likely to have lifetime PTSD than men, even when risk of exposure and types of trauma are taken into consideration^3, 4, 8, 10–12^. This sex bias in disease prevalence is also observed in depression^13^ and other commonly occurring mood and anxiety disorders^14^ that are heavily influenced by stress exposure. Preclinical and clinical investigations have identified sexual dimorphism in stress response systems that may be involved in the increased prevalence of stress-related psychopathologies in women^15, 16^. Furthermore, in addition to sexual dimorphism in the neurobiological underpinnings of stress/trauma response, recent animal studies suggest that behavioral response to traumatic stress is fundamentally different between males and females and should be considered in interpretation of results^17^. In humans, response to stress/trauma exposure involves both biological and social contributors corresponding to sex and gender-related variables. While the effects of sex and gender are difficult to disentangle, investigations stratified by biological sex, understood to interact with gender-related variables, are warranted to improve our currently limited understanding of the sex-specific biological processes dysregulated in PTSD pathophysiology.

Mounting evidence suggests a key role for stress-induced inflammation and immune alteration in the development and maintenance of PTSD and other stress-related psychiatric disorders. Although findings from human literature have been mixed, PTSD has generally been associated with increased pro-inflammatory tone, basally and in response to immune challenge, via both cytokine signaling and changes in immune cell distribution/function^18–34^. Investigations in animal models, specifically male rodent studies, have provided mechanistic insights into how peripheral immune cell response/signaling and distribution is linked with microglial activation and neuroinflammatory dynamics to trigger stress-induced anxiety behavior and memory impairment^35–40^.

Epigenetic mechanisms have emerged as important regulators of PTSD-associated immune dysregulation and inflammation^41–46^, and are particularly significant for the study of PTSD because they capture the interactions among pre-disposing genetic/environmental risk factors and the precipitating trauma exposure. Individual response to trauma exposure can be encoded as short-lived or persistent epigenetic changes that reflect and may contribute to posttraumatic physiological changes, some of which remain after remission of PTSD symptoms. DNA methylation (DNAm) has been the most widely studied epigenetic mechanism and evidence from both animal and human models point to its key role in stress regulation^9, 47–49^, fear memory^50–54^, and immune function^43–46, 55–58^, in both brain and blood. Exploring PTSD-associated DNAm profiles in blood may inform our understanding of mechanisms underlying immune dysregulation, particularly those that coordinate peripheral immune-neuroimmune crosstalk^59, 60^.

Peripheral blood-based methylomic profiles are comprised of two dynamic components: 1) profiles reflecting proportions of immune cell subtypes (i.e., leukocyte composition) and 2) alterations in DNAm levels at CpG sites genome-wide (i.e., differential methylation). Epigenome-wide association studies (EWAS) often seek to identify dynamic differential methylation marks and treat cellular heterogeneity as a major confound that must be addressed to improve signal detection. However, shifts in leukocyte subtypes provide key insights into immunological changes and warrant examination themselves. Recent developments in deconvolution algorithms and cell-type discriminating reference databases have improved estimates^61–64^ and enabled utility of DNAm-based leukocyte subtype estimates as proxies for white blood cell differential-based metrics^65, 66^. Leveraging these recently developed methods, here we use leukocyte-derived methylomic profiles combined from two civilian cohorts to investigate our hypothesis that PTSD is associated with sex-specific shifts in leukocyte composition, detectable by DNAm-based estimates. To our knowledge, this study is the first to investigate leukocyte composition profiles in PTSD using these new DNAm-based approaches for immune profiling.

## 2. MATERIALS AND METHODS

### 2.1. Study Participants

Trauma-exposed, adult participants with Illumina HumanMethylation450 (450K) BeadChip array data were selected from two predominantly African-American, community-based cohorts examining biological and environmental correlates of PTSD, namely the Detroit Neighborhood Health Study (DNHS) and Grady Trauma Project (GTP). The DNHS, based in Detroit, MI, was approved by the institutional review boards of the University of Michigan and University of North Carolina at Chapel Hill. The GTP, based in Atlanta, GA, was approved by the institutional review boards of Emory University School of Medicine and Grady Memorial Hospital. All participants provided written informed consent prior to data collection. Details regarding the DNHS^43, 67, 68^ and GTP^69–71^ were published previously. While neither study excluded participants based on illness, women known to be pregnant (in the GTP) were excluded from estimation and analyses, due to well-known/known significant differences in leukocyte composition during pregnancy^72^. Collected demographic data included self-reported gender, race, age, and current smoking, which was defined as any cigarette smoking in the past 30 days.

### 2.2 Assessment of PTSD

Study participants were assessed for PTSD, as defined by the Diagnostic and Statistical Manual of Mental Disorders, Fourth Edition (DSM-IV)^1^. In the DNHS, study participants were assessed for PTSD using the well-validated self-report PTSD Checklist, Civilian Version (PCL-C)^73–76^ and additional questions about duration, timing, and impairment due to symptoms, via structured telephone interviews^10, 43, 77^. Participants who met all six DSM-IV criteria in reference to their worst traumatic event or to a randomly selected traumatic event (if the participant experienced more than one trauma), were considered affected by lifetime PTSD. Those that met criteria based on symptoms reported within the past month were considered affected by current PTSD. The survey-based diagnoses were validated in a random subsample of participants via in-person clinical interview and showed high internal consistency and concordance^43, 78^. In the GTP, study participants were assessed for lifetime and current PTSD using the clinician-administered PTSD scale (CAPS)^79^.

### 2.3 Sample Processing

Genomic DNA was extracted from peripheral blood using the DNA Mini Kit (Qiagen, Germantown, MD) in the DNHS and the Gentra Puregene Kit (Qiagen, Germantown, MD) in the GTP. Both studies bisulfite-converted the DNA samples using the Zymo EZ-96 DNA methylation kit (Zymo Research, Irvine, CA) and whole-genome amplified, fragmented, and hybridized the samples to the Illumina Human Methylation 450K BeadChip array (Illumina, San Diego, CA), according to the manufacturers’ recommended protocols, as published previously^58,71,80–82^.

### 2.4 Quality control and pre-processing of 450K Data

The raw .idat files were imported into R (version 3.5.1)^83^, using the *minfi*^84^ Bioconductor (version 3.7)^85, 86^ package, for all subsequent data processing and analyses. After quality control (QC)^84, 87, 88^, data pre-processing^89–94^ was conducted on all QC’ed samples (DNHS: n = 187; GTP: n = 390). This included duplicates in the DNHS and participants with missing data in the GTP. Analyses were conducted on unique participants that passed QC, as described below, and had PTSD data available (Table 1).

**Table 1:** Key demographic characteristics of the DNHS and GTP

|  | **DNHS**<br>**(n = 175)** | **GTP**<br>**(n = 308)** | **Total**<br>**(n = 483)** |
| --- | --- | --- | --- |
| **Sex** |  |  |  |
| Female | 111 (63.4%) | 219 (71.1%) | 330 (68.3%) |
| Male | 64 (36.6%) | 89 (28.9%) | 153 (31.7%) |
| **Race** |  |  |  |
| African American | 149 (85.1%) | 282 (91.6%) | 431 (89.2%) |
| Caucasian | 23 (13.1%) | 21 (6.8%) | 44 (9.1%) |
| Other | 3 (1.7%) | 5 (1.6%) | 8 (1.7%) |
| **Median Age (IQR)** |  |  |  |
|  | 54 (45.5-62.5) | 45 (33-52) | 48 (37.5-55) |
| **Current Smoking** |  |  |  |
| No | 106 (60.6%) | 176 (57.1%) | 282 (58.4%) |
| Yes | 68 (38.9%) | 119 (38.6%) | 187 (38.7%) |
| Missing | 1 (0.6%) | 13 (4.2%) | 14 (2.9%) |
| **Lifetime PTSD** |  |  |  |
| No PTSD | 59 (33.7%) | 147 (47.7%) | 206 (42.7%) |
| PTSD | 116 (66.3%) | 161 (52.3%) | 277 (57.3%) |

This table describes the subset of participants included in primary analyses investigating sex-specific associations between DNAm-based cell estimates and lifetime PTSD. For the 2-way ANCOVA, 14 participants were excluded due to missing current smoking data.
DNHS: Detroit Neighborhood Health Study; GTP: Grady Trauma Project

For data quality assessment, samples were checked for 1) low total signal (mean signal intensity less than half of the overall median, after setting probes with detection p-value > 0.001 or < 2,000 arbitrary units to missing); 2) > 1% of failed probes (detection p-value > 0.001); 3) outlying beta value distribution (i.e., smaller than three times interquartile range (IQR) from the lower quartile or larger than 3 times IQR from the upper quartile); 4) greater than three standard deviations of the mean bisulfite conversion control probe signal intensity^64^. Additionally, samples were checked for gender discordance based on median total intensity of X and Y-chromosome mapped probes (as implemented in *minfi*^84^) and removed if predicted sex differed from self-reported gender. Five samples were removed among DNHS samples for gender discordance, and six samples were removed among GTP samples (two for data quality and four for gender discordance). After within-array background correction and dye-bias equalization using out-of-band control probes (ssNoob^90, 95^; *minfi*), probes with detection p-value > 0.001 in more than 10% of samples^87^ and cross-reactive probes^91^ (i.e., cross-hybridized between autosomes and sex chromosomes) were removed. Beta-mixture quantile (BMIQ) normalization (*ChAMP*^93, 94^) was used to correct for type II probe bias^92^.

To control for technical artifacts (e.g., sample processing and imaging batch effects), PCs based on non-negative control probe signal intensity^88^ were removed from BMIQ-normalized M-values (i.e., logit-transformed beta-values) separately for each study. PC correlation heatmaps were used to check for successful removal/reduction of batch effects, especially chip and row effects, while maintaining signal from biological variables. The DNHS and GTP datasets were then combined and an empirical Bayes method (i.e., ComBat^96^ in the *sva* package^97^) was used on the combined M-values to account for study effects while controlling for sex and age. Data quality assessment, QC probe filtering, and first step of batch removal were study-specific, while pre-processing steps (ssNoob+BMIQ) implemented within-array approaches unaffected by study. Only probes that passed QC in both studies (n = 455,072 probes) were included in the combined dataset.

### 2.5 Leukocyte Composition Estimation

Leukocyte composition was estimated on ComBat-adjusted beta-values using the *EpiDISH*^62^ reference database, which is informed by cell-type specific DNase hypersensitive sites (DHS; based on the NIH Epigenomics Roadmap database^98^) and is optimized for discriminating granulocytes, CD14^+^ monocytes, CD8^+^ T cells, CD4^+^ T cells, CD19^+^ B cells, and CD56^+^ natural killer cells. Robust partial correlation (RPC; robust multivariate linear regression, non-constrained projection) was used as the primary deconvolution algorithm and *EpiDISH’s* implementation of linear, constrained projection (CP), originally introduced by Houseman et al. (2012)^99^, was used to calculate a second set of estimates for comparison.

### 2.6 Ancestry Estimation

DNAm-based ancestry principal components (PCs) were derived on cleaned beta-values after regressing out sex and age from batch-adjusted M-values. Ancestry PCs were calculated on a subset of 2,317 ancestry informative CpG probes included in two published ancestry informative CpG lists that accounted for confounders^100^ and that included probes within 10 base pairs (bp) of single nucleotide polymorphisms (SNPs)^101^. The first 2 PCs based on this subset of probes were used as ancestry PCs (ancPCs) after checking for strong association with self-reported race and effective separation of self-reported races.

### 2.7 Statistical Analysis

The Shapiro-Wilk test was used to assess normality and Levene’s test was used to compare equality of variance among groups. Since cell estimates had dissimilar, non-normal distributions, non-parametric tests were used for initial group comparisons of leukocyte subtypes. The two-sample Kolmogorov-Smirov (KS) test was used to compare distribution of cell estimates when variances were unequal between groups and Mann-Whitney U test was used to compare mean ranks of cell estimates otherwise. Spearman’s rank correlation was used to assess agreement between estimates based on RPC and CP deconvolution approaches. A threshold of 0.05 was used for p-values and p-values were adjusted for multiple comparisons using Holm’s method (citation), unless otherwise specified.

To test our main hypothesis—that PTSD is associated with sex-specific shifts in leukocyte composition—initial sex-stratified analyses were conducted on all leukocyte subtypes using the non-parametric Mann-Whitney U test. For leukocyte subtypes determined to be significantly associated with lifetime PTSD in either sex based on initial Mann-Whitney U tests, a two-way analysis of covariance (ANCOVA; Type III) controlling for age, ancestry (based on DNAm ancestry PCs), and current smoking, was performed with post-hoc comparison of estimated marginal means to examine the effects of sex and lifetime PTSD on transformed cell estimates. Transformation for cell estimate was informed by conducting Tukey’s Ladder of Powers to maximize normality based on the Shapiro-Wilks W statistic. Sex-stratified Kruskal-Wallis and post-hoc Dunn tests were conducted as additional follow-up to investigate possible differences in cell proportions by PTSD status (i.e., trauma-exposed controls, remitted PTSD, and current PTSD).

## 3. RESULTS

### 3.1 Demographic characteristics of sampled study participants from the DNHS and GTP

The demographic characteristics of study participants included in primary analyses investigating sex-specific associations between DNAm-based cell estimates and lifetime PTSD are presented in Table 1. Of the 483 participants from the combined DNHS and GTP sample, 57.3% had a lifetime diagnosis of PTSD, 68.3% were female, and 38.7% were current smokers. The study population was predominantly African-American (89.2%), based on self-reported race, and the median age was 48 years (IQR: 17.5; 37.5-55 years).

### 3.2 Comparison of leukocyte subtype estimates by sex and lifetime PTSD

Cell estimates were compared by sex and lifetime PTSD in each leukocyte subtype to establish overall differences. Significant overall sex differences were observed in the distributions of CD56^+^ natural killer (NK) cell (KS: *D* = 0.19, *adj. p* = 0.007) and CD8^+^ T cell (CD8T) proportions (KS: *D* = 0.16; *adj. p* = 0.04) in RPC estimates. Males showed greater variability than females for both NK and CD8T cells (male vs. female - IQR_NK_: 6.2% vs. 4.15%; IQR_CD8T_: 9.5% vs. 6.2%), as well as higher median NK (5.5% vs. 4.4%) and lower median CD8T (9.0% vs. 9.7%) cell proportions (Figure 1). Significant sex differences were also observed in the comparison set of CP estimates for the distributions of both NK (KS: *D* = 0.21, *adj p* = 0.001) and CD8T (KS: *D* = 0.25, *adj p* = 4.21×10^−5^) cell estimates, with males again showing greater variability than females for both NK and CD8T cells (male vs. female - IQR_NK_: 7.3% vs. 5.2%; IQR_CD8T_: 6.4% vs. 5.1%), as well as higher median NK (5.0% vs. 2.9%) and lower median CD8T (11.6% vs. 13.5%) cell proportions (Supplementary Figure 1). No significant overall differences (i.e., in both sexes combined) were observed between lifetime PTSD cases and trauma-exposed controls in any leukocyte subtype, based on either RPC or CP estimate (Levene’s test and Mann-Whitney U test).

**Figure 1:**
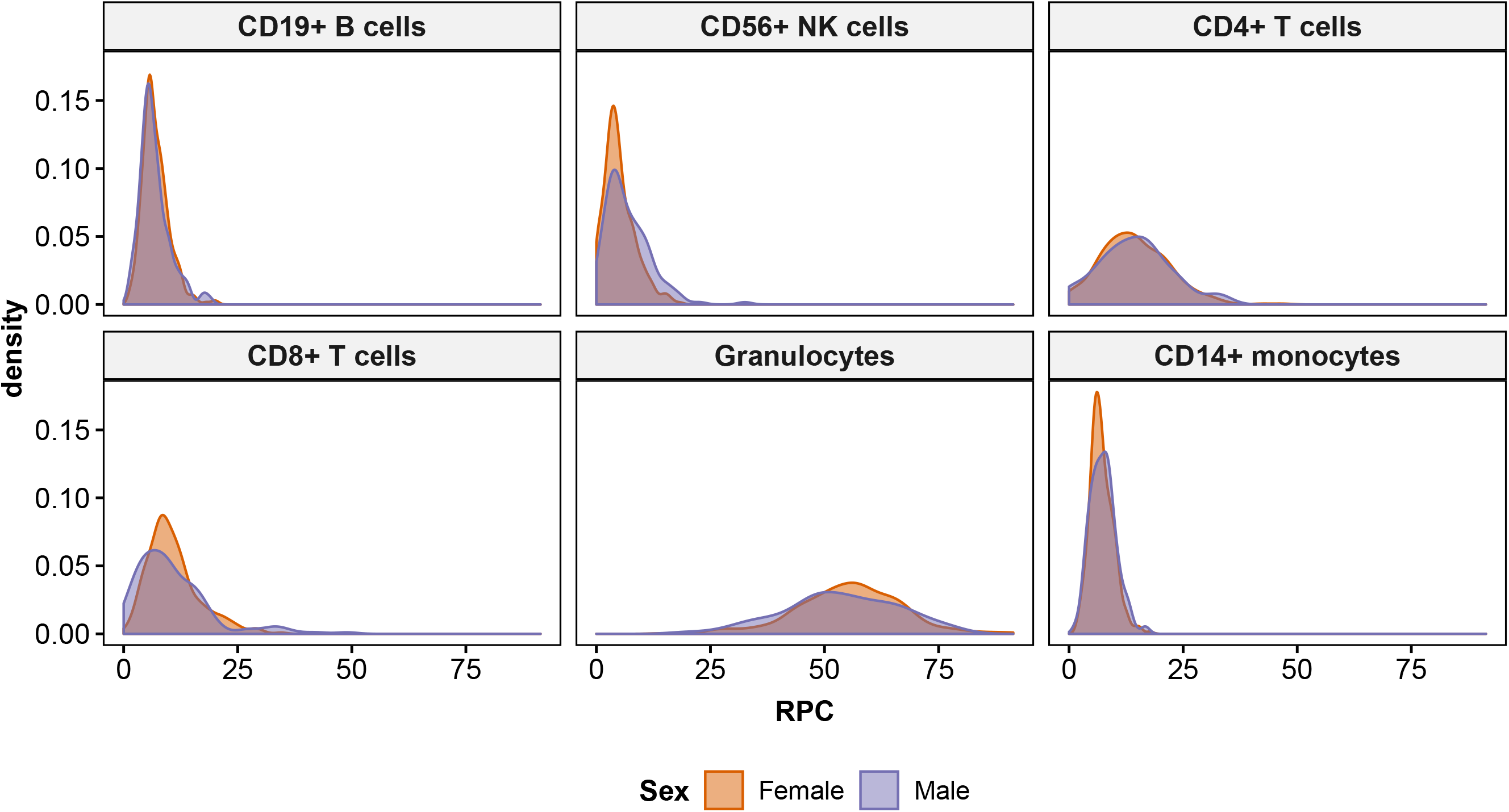
Distribution of leukocyte subtypes based on robust partial correlation (RPC) estimates, by sex. Sex differences in CD8^+^ T and CD56^+^ NK cell distributions were found to be prominent.

### 3.3 Comparison of leukocyte subtype estimates by deconvolution approach

Good overall agreement was observed between RPC and CP estimates, as measured by Spearman’s correlation (i.e., RPC-CP correlation), but CD8T cells showed much poorer agreement, *ρ_s_*(481) = 0.83, relative to the other leukocyte subtypes, *ρ_s_*(481) > 0.94. Since the main objective of this study was to investigate sex-specific shifts in leukocyte composition, comparison of RPC and CP estimates was stratified by sex (Supplementary Figure 2). Sex-stratified RPC-CP correlation revealed that the largest difference in RPC-CP correlation between sexes was also found in CD8T cells, |Δ*ρ_s_*| = 0.07, such that females showed poorer correlation, *ρ_s_*(328) = 0.80, than males, *ρ_s_*(151) = 0.87. For the other leukocyte subtypes, the difference in RPC-CP correlation between sexes (|Δ*ρ_s_*|), ranged from 0.01 to 0.03, with CD56^+^ natural killer (NK) cells having the second largest difference in correlation between sexes (female: *ρ_s_*(328) = 0.93; male: *ρ_s_*(151) = 0.96). In all leukocyte subtypes, except CD19^+^ B cells, females had lower correlation coefficients than males.

### 3.4 Elevated monocyte proportions are associated with lifetime PTSD in males, but not females

Sex-stratified Mann-Whitney U tests revealed a significant difference in monocyte proportions between PTSD cases and controls in males (Figures 2 and 3). Males with lifetime PTSD had higher median monocyte proportions than trauma-exposed controls, *U* = 2100, *Z* = −2.9, *p* = 0.004, *adj. p* = 0.026, *r* = 0.23. In contrast, no difference in monocyte estimates was found between groups in females, *U* = 13000, *Z* = −0.58, *p* = 0.6, *adj. p* = 1, *r* = 0.03. Lifetime PTSD-associated differences were not observed in any other leukocyte subtypes in either sex.

**Figure 2:**
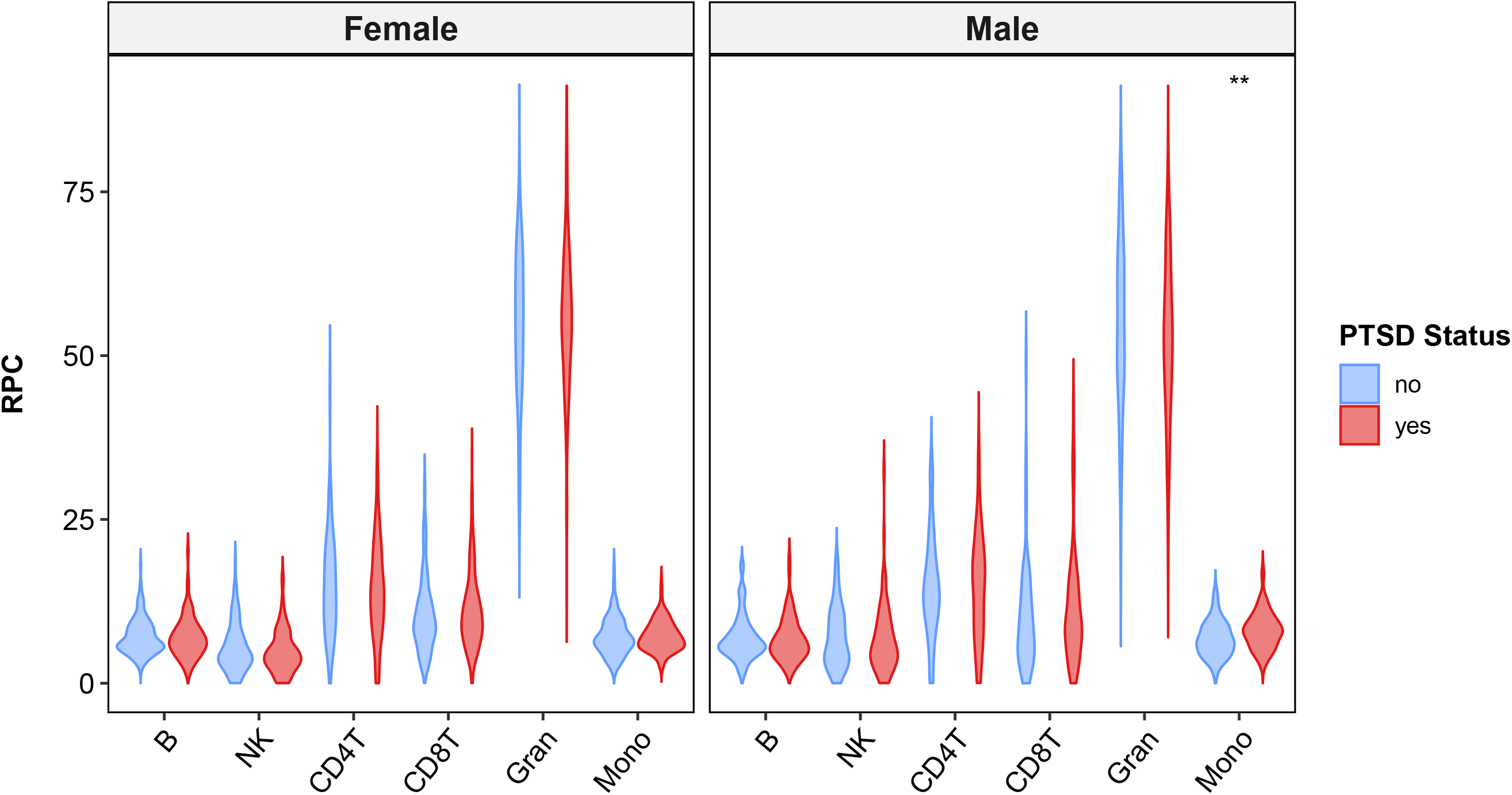
Violin plots of RPC estimates for lifetime PTSD cases and controls, stratified by sex. Only monocyte proportions in males showed statistically significant difference between lifetime PTSD cases and controls, based on Mann-Whitney U test (p-value = 0.004; Holm-adjusted p-value = 0.026). For figure labels on x-axis: B = CD19^+^ B cells; NK = CD56^+^ NK cells; CD4T = CD4^+^ T cells; CD8T = CD8^+^ T cells; Gran = Granulocytes; Mono = CD14^+^ monocytes.

**Figure 3:**
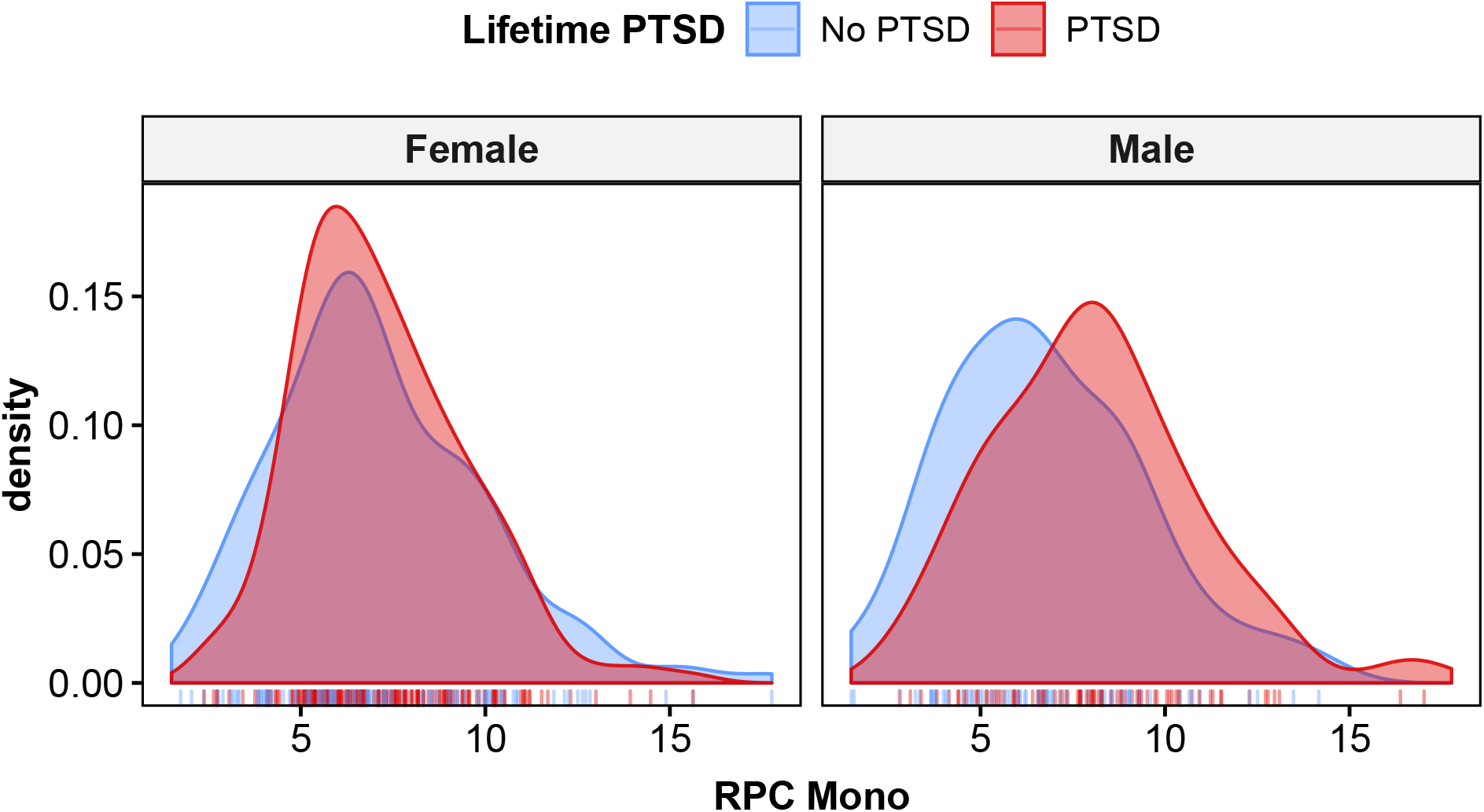
Density plots for RPC monocyte estimates in lifetime PTSD cases and controls, stratified by sex, show distinctly higher monocyte levels in males with lifetime PTSD compared to trauma-exposed cases. This difference in monocyte levels between lifetime PTSD case and controls is not observed in females.

A two-way ANCOVA was conducted to investigate whether sex moderated the effects of lifetime PTSD on square root transformed monocyte estimates, while accounting for age, ancestry, and current smoking (Table 2). A significant interaction was found between sex and lifetime PTSD, *F*(1, 461) = 4.89, *p* = 0.027, η_p_^2^ = 0.011. Post-hoc comparison of estimated marginal means (EMMs) for lifetime PTSD by sex (Figure 4; Table 3) showed a significant mean difference between lifetime PTSD cases and controls in males, Δ*EMM* = 0.26, *SE* = 0.08, *t*(461) = 3.32, *p* = 0.001, where mean monocyte estimates were higher in lifetime PTSD cases than controls. No significant mean difference was observed between PTSD cases and controls in females, Δ*EMM* = 0.05, *SE* = 0.05, *t*(461) = 0.89, *p* = 0.37, confirming findings from initial sex-stratified analyses. Together, our results suggest that male PTSD cases have significantly elevated monocyte proportions compared to trauma-exposed controls and that this lifetime PTSD-associated shift is not observed in females.

**Figure 4:**
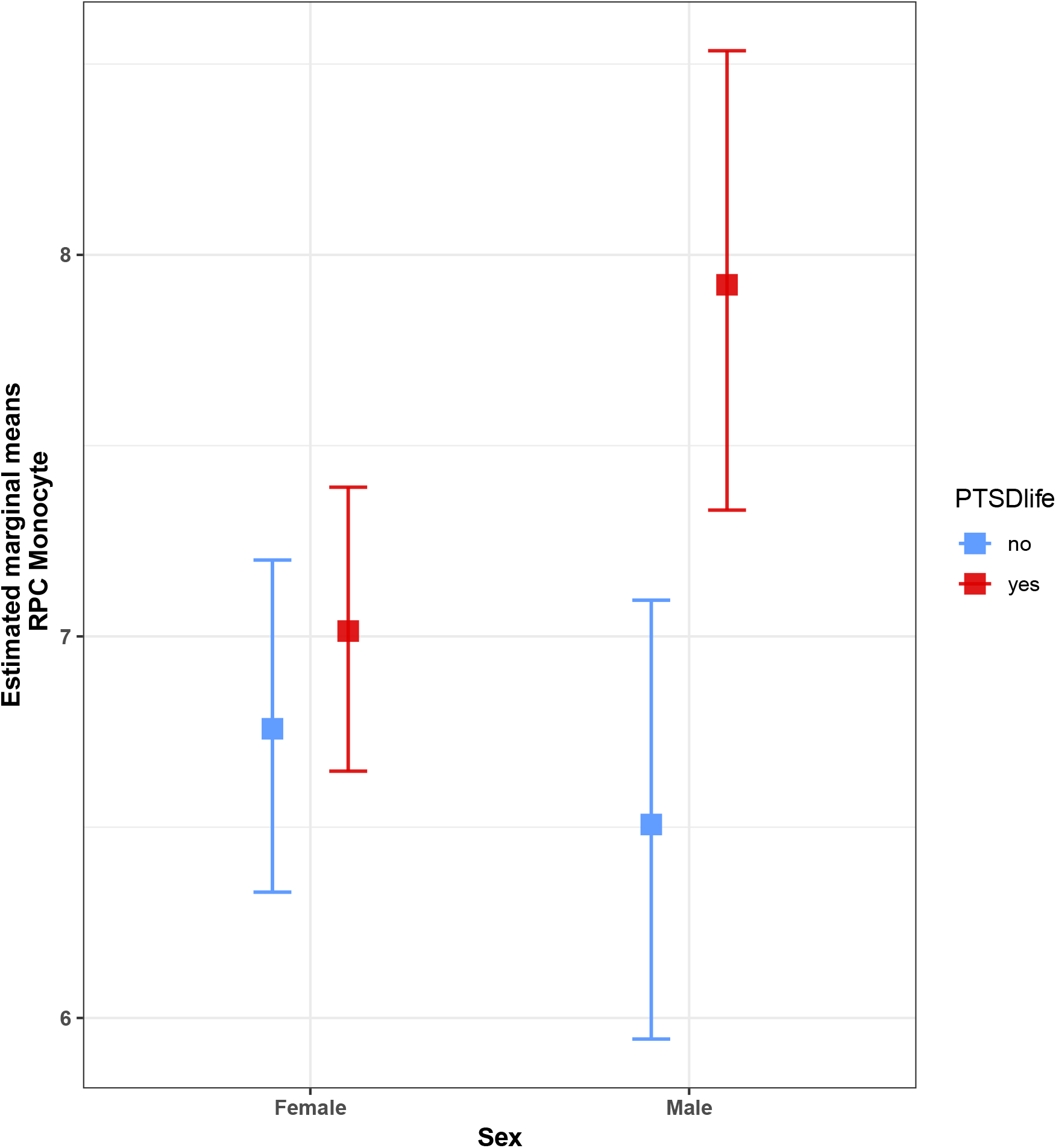
Lifetime PTSD by sex interaction plot for estimated marginal means (EMMs) of RPC monocyte estimates. Interaction plot shows a significant EMM difference between lifetime PTSD cases (red) and controls (blue) in males, where mean monocyte estimates are higher in cases than controls. No significant EMM difference was observed between PTSD cases and controls in females.

**Table 2:** Two-way ANCOVA Table for RPC monocyte estimates (n = 469)

| Terms | Type III Sum of Squares | *df* | Mean Square | *F* | *p* | partial $\eta^2$ |
| --- | --- | --- | --- | --- | --- | --- |
| Sex | 0.336 | 1 | 0.336 | 1.469 | 0.226 | 0.003 |
| PTSDlife | 2.379 | 1 | 2.379 | 10.407 | 0.001*** | 0.022 |
| Age | 2.309 | 1 | 2.309 | 10.101 | 0.002*** | 0.021 |
| ancPC1 | 0.000 | 1 | 0.000 | 0.001 | 0.977 | 0.000 |
| ancPC2 | 0.219 | 1 | 0.219 | 0.957 | 0.329 | 0.002 |
| Smoking | 0.060 | 1 | 0.060 | 0.261 | 0.610 | 0.001 |
| Sex:PTSDlife | 1.146 | 1 | 1.146 | 5.011 | 0.026* | 0.011 |
| Residuals | 105.399 | 461 | 0.229 |  |  |  |

\*\* $p < 0.05$ , \* $p < 0.01$ , \*\*\* $p < 0.005$

**Table 3:** Summary for RPC monocyte estimates by group

| **Sex** | **PTSD** | **n** | **mean** | **SE** | **EMM** | **SE<sub>EMM</sub>** | **lower.CL** | **upper.CL** |
| --- | --- | --- | --- | --- | --- | --- | --- | --- |
| Female | no | 135 | 7.113 | 0.2443 | 6.758 | 0.2214 | 6.330 | 7.200 |
| Female | yes | 184 | 7.182 | 0.1691 | 7.014 | 0.1893 | 6.647 | 7.391 |
| Male | no | 70 | 6.803 | 0.3228 | 6.507 | 0.2926 | 5.945 | 7.095 |
| Male | yes | 80 | 8.103 | 0.3147 | 7.921 | 0.3063 | 7.331 | 8.535 |

This table describes untransformed RPC monocyte estimates by group (i.e., sex and lifetime PTSD); n = count per group; EMM = estimated marginal means (i.e., least squares means); SE = standard errors for regular means; SE<sub>EMM</sub> = standard errors for EMM.
Lower and upper confidence limits are for 95% level. EMM and intervals were back-transformed from the square root scale to the original scale of cell subtype proportions (%).
Significance level of alpha = 0.05 was used for EMM comparisons. Results for pairwise comparison were averaged over levels for current smoking. Degree of freedom was 461 and male lifetime PTSD cases were significantly different from other groups.

### 3.5 Association between monocyte proportions and lifetime PTSD in males is independent of current PTSD status

To investigate whether participants with current PTSD exhibited a different monocyte profile from those with remitted PTSD, a sex-stratified Kruskal-Wallis test was conducted for PTSD status (i.e., trauma-exposed controls, remitted PTSD, and current PTSD; Figure 5). A significant difference in monocyte estimates was observed in males, *H*(2) = 8.2, *p* = 0.017, but not in females, *H*(2) = 1.1, *p* = 0.59, confirming findings from analyses for lifetime PTSD. The post-hoc Dunn test revealed significant differences between PTSD case groups and trauma-exposed controls (current PTSD vs. controls: *Z* = 2.31, *p* = 0.021, *adj. p* = 0.042, *r* = 0.23; remitted PTSD vs controls: *Z* = 2.40, *p* = 0.016, *adj. p* = 0.049, *r* = 0.22), but no significant difference between current and remitted PTSD groups, *Z* = 0.18, *p* = 0.86, *adj. p* = 0.86, *r* = 0.02. These findings suggest that the association between monocyte proportions and lifetime PTSD in males is independent of current PTSD state and may reflect long-standing changes associated with lifetime history of PTSD diagnosis.

**Figure 5:**
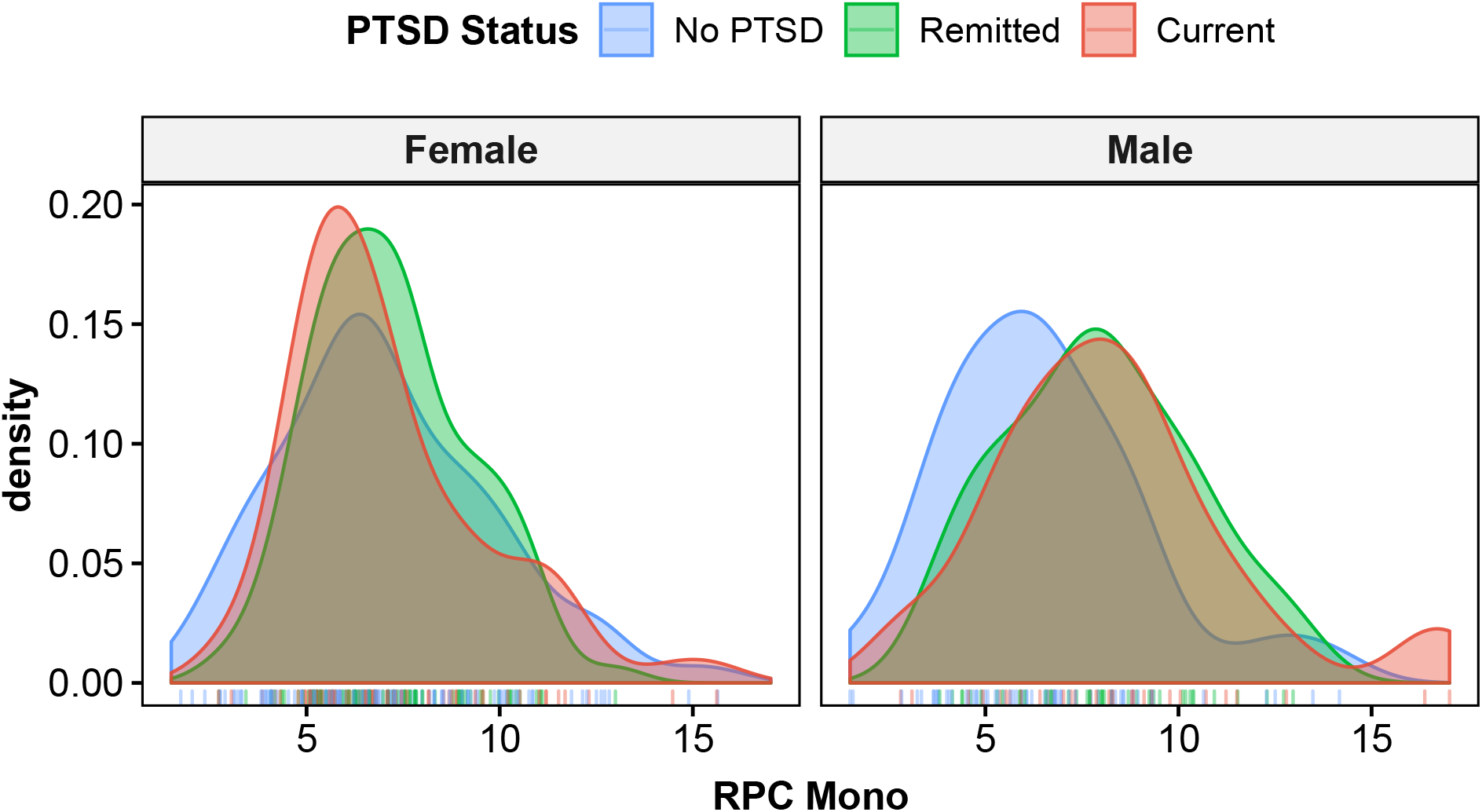
Density plots for RPC monocyte estimates comparing those with current PTSD, remitted PTSD, and trauma-exposed controls, stratified by sex. Distinguishing between current and remitted PTSD cases suggests that the significant peak shift in male PTSD cases is associated with long-standing PTSD trait, rather than current PTSD state. Corresponding post-hoc Dunn test revealed no significant difference between current and remitted PTSD cases and significant differences between PTSD case groups and trauma-exposed controls. Again, no significant differences were observed in females.

### 3.6 CP-based estimates confirm male-specific association between lifetime PTSD and DNAm-based monocyte proportions

Comparative analyses using CP-based monocyte estimates confirmed our significant sex-specific finding between lifetime PTSD and monocyte proportions. As observed previously, males with lifetime PTSD had higher monocyte proportions than trauma-exposed controls, *U* = 2200, *Z* = −2.5, *p* = 0.01, *r* = 0.20, while no difference was found between PTSD cases and controls in females, *U* = 13000, *Z* = −0.15, *p* = 0.88, *r* = 0.008 (Supplementary Figure 3). A two-way ANCOVA on square root transformed CP-based estimates, accounting for age, ancestry, and current smoking (Supplementary Table 1), showed a significant interaction between sex and lifetime PTSD, *F*(1,461) = 3.99, *p* = 0.046, η_p_^2^ = 0.009. Post-hoc comparison of EMMs by sex (Supplementary Figure 4; Supplementary Table 2) confirmed a significant mean difference between lifetime PTSD cases and controls in males, with higher mean monocyte estimates in PTSD cases, Δ*EMM* = 0.24, *SE* = 0.09, *t*(461) = 2.85, *p* = 0.005. Again, no significant mean difference between PTSD cases and controls was found in females, Δ*EMM* = 0.04, *SE* = 0.06, *t*(461) = 0.63, *p* = 0.53. Smaller difference in EMM (Δ*EMM*) and larger EMM standard error were observed in male CP estimates, compared to RPC estimates.

Finally, sex-stratified Kruskal-Wallis tests for PTSD status using CP-based monocyte estimates (Supplementary Figure 5) also suggested significant differences in males, *H*(2) = 6.5, *p* = 0.038, but not females, *H*(2) = 0.74, *p* = 0.69. While no difference was observed between current and remitted PTSD groups, *Z* = 0.24, *p* = 0.81, *adj. p* = 0.81, *r* = 0.03, in the post-hoc Dunn test, differences between PTSD cases and trauma-exposed controls showed smaller effect sizes and only nominal significance (current PTSD vs controls: *Z* = 2.11, *p* = 0.035, *adj. p* = 0.10, *r* = 0.21; remitted PTSD vs controls: *Z* = 2.10, *p* = 0.036, *adj. p* = 0.071, *r* = 0.19).

Overall, confirmation of primary findings in RPC-based monocyte estimates using CP estimates suggests that male-specific association of DNAm-based monocyte proportions with lifetime PTSD is consistent across deconvolution approaches. This is supported by lifetime PTSD analyses (i.e., Mann-Whitney U test, two-way ANCOVA, and post-hoc comparison of EMMs) and by comparison between current and remitted PTSD groups in males, which revealed no significant difference (post-hoc Dunn test). However, CP-based results consistently presented smaller effect sizes than RPC-based results across all analyses and follow-up comparisons between PTSD case groups and trauma-exposed controls did not reach significance after p-value adjustment in post-hoc Dunn test using CP monocyte estimates in males.

## 4. DISCUSSION

The primary goal of the present study was to test whether PTSD is associated with sex-specific shifts in leukocyte composition, detectable by DNAm-based estimates; in addition, we sought to implement two leukocyte composition estimation approaches to assess consistency of results across different deconvolution algorithms. We found that lifetime PTSD was associated with a small but significant elevation in monocyte proportions in males, but not females. No difference in monocyte proportions was observed between current and remitted PTSD cases in males, suggesting that this sex-specific peak shift reflects a long-standing trait of lifetime history of PTSD diagnosis, rather than current state of PTSD. Additionally, these findings were observed in both the primary robust partial correlation (RPC) and comparison constrained projection (CP)-based sets of cell estimates, which were derived using non-constrained vs. constrained projection deconvolution algorithms, respectively. Overall, our main finding of elevated monocyte proportions in males, but not females with lifetime history of PTSD provides evidence for a sex-specific shift in peripheral blood leukocyte composition that is detectable in methylomic profiles and that reflects long-standing changes associated with PTSD diagnosis.

Methylomic profiles derived from peripheral blood offer a wealth of information on immune state that can be harnessed to detect shifts in leukocyte composition and epigenetic changes in regulation of immune processes. Methylomic profiles are particularly well-suited for investigating PTSD because DNAm encodes individual response to trauma and may play a key role in PTSD-associated immune dysregulation. Our study leveraged recent advances in reference-based deconvolution methods^62–64^ to investigate sex-specific shifts in leukocyte composition associated with PTSD. Specifically, we implemented the *EpiDISH* algorithm^62^, which (i) employs DNase hypersensitive site (DHS) data of leukocyte subtypes to inform probe selection for their reference database and (ii) introduces robust partial correlation (RPC), a non-constrained projection approach for reference-based deconvolution. Previous comparisons across deconvolution approaches suggest that RPC may, in general, be more robust than the established constrained projection (CP) approach^99^ for realistic noise levels^62, 64^. Although our main finding in RPC-based monocyte estimates was confirmed using CP, effect sizes were consistently smaller and EMM standard errors were larger in CP-based monocyte estimates. Additionally, follow-up comparisons between PTSD case groups and trauma-exposed controls in males did not reach significance after p-value adjustment in CP-based post-hoc Dunn test, despite significant results in other analyses of lifetime PTSD. Thus, RPC-based estimates were better able to resolve male-specific association of monocyte proportions with lifetime PTSD consistently across all conducted analyses compared to CP-based estimates and our results also favor use of RPC estimates for modeling leukocyte composition.

Comparison between RPC and CP estimates, based on the same *EpiDISH* reference database (of 330 leukocyte subtype discriminating CpG probes), further illustrates contexts where choice of deconvolution approach matters. Since RPC applies the non-negative and normalization constraints after deriving coefficients/proportion estimates and models by robust regression^62^, it is likely to demonstrate advantages over CP more clearly when the reference database does not adequately identify or account for major functional subsets of leukocyte subtypes (e.g., naïve vs. effector vs. memory CD8T cells). This may have been the case for CD8T cells, which exhibited much poorer correlation between RPC and CP estimates than other leukocyte subtypes and the largest difference in RPC-CP correlation between sexes. Probe selection for CD8T cells may be challenged by the active role of DNAm dynamics in regulating key processes that define CD8T cells^102^ and sex differences in RPC-CP correlation may reflect the poorer fit to CD8T discriminating probes during CP estimation in females. Further investigation of baseline sex differences in cell-specific DNAm-levels, especially at leukocyte subtype-discriminating CpG probes (i.e., reference database), is warranted.

Comparison of leukocyte subtype estimates by sex revealed significant baseline sex differences in the distributions of NK cell and CD8T cell proportions, with males showing greater median NK and lower median CD8T cell proportions, using both RPC and CP based estimates. This finding is consistent with a previous study that reported sex differences in both leukocyte subtypes using estimates based on *minfi’s* implementation of the Houseman approach^103^ and with immunology literature that reported higher NK cell counts and proportions in males compared to females^104^. A recent study that modeled cell-specific methylation profiles also reported robust sex differences in CD56^+^ NK methylation patterns^105^, suggesting that this leukocyte subtype may be regulated by DNAm in a sex-specific manner. Additionally, DNAm dynamics have been found to drive effector functions in CD8T cells after stimulation^102, 106^. Development of reference databases that resolve the six main leukocyte subtypes to consider proportions of subsets with shared lineage but different functionality/phenotype (e.g., naïve vs memory vs regulatory subtypes) may allow us to explore this hypothesis and would greatly enrich our understanding of immune activity.

Our main finding of elevated monocyte proportions in male lifetime PTSD cases is consistent with previously reported evidence. While not specifically in PTSD, a previous study of Gulf War Illness (GWI) in a predominantly male veteran cohort reported an association between GWI and increase in monocyte count^107^. Further prospective investigation in humans is needed to determine whether this male-specific difference in monocyte proportions reflects an increased susceptibility for developing PTSD or if it reflects an immunological shift in response to the precipitating trauma associated with PTSD psychopathology. Studies in male rodent models provide strong evidence for the latter and have been important for establishing the relationship between peripheral immune cells and the brain in the context of psychosocial stress and associated behavior. For example, repeated social defeat (RSD) was found to induce myelopoiesis and release inflammatory monocytes into circulation via sympathetic signaling, and this increased level of circulating peripheral monocytes was associated with recruitment of pro-inflammatory monocytes/macrophages to the brain and neuroinflammation^35, 108, 109^. Macrophage recruitment to the brain was demonstrated to correspond with development, maintenance, and re-establishment of RSD-induced anxiety-like behavior, and blockade of this recruitment (via splenectomy or β-adrenergic receptor blockage) before RSD was found to prevent development of anxiety-like behavior^37, 38^. Additionally, a recent paper discerned that stress-induced anxiety-like behavior and social avoidance are dependent on an increase in IL-6 after stress exposure, which induces a primed transcriptional profile in monocytes recruited to the brain and propagates IL-1 mediated inflammation associated with anxiety-like behavior^110^. These studies implicate peripheral monocytes in directly affecting PTSD-like behavior after stress exposure in males.

Unfortunately, all the aforementioned studies have been in males, so it is not clear to what extent these mechanistic insights pertain to females^111, 112^ and recent investigations in rodent models suggest fundamental sex differences in neurobiological response to trauma exposure^17^. Chronic PTSD-associated sex differences were noted in transcriptional regulation^113^ and gene expression^114^ of CD14^+^ monocytes isolated from peripheral blood, however no studies of PTSD have investigated sex differences in monocyte methylomic profiles. Given the inherent sex differences in innate immune response^115, 116^, as understood in the context of infection, injury, and treatment of inflammatory disorders, sexually dimorphic dynamics and effects may also exist in the context of neuroimmune response to stress/trauma exposure^111^. A study examining sex differences in regulation of inflammatory cell recruitment and cytokine synthesis found that ovarian hormones regulate phenotype, function, and numbers of macrophages, but not T lymphocytes, in females^117^. This fundamental sex difference may underlie more efficient recognition and elimination of infectious stimuli without recruitment of circulating neutrophils or excessive cytokine production in females, compared to males^117^, and may also have implications in the context of psychosocial stress exposure. Our observation of male-specific increase in monocyte proportions associated with lifetime PTSD may reflect fundamental sex differences in leukocyte trafficking, tissue distribution, and thus composition in blood, with implications for stress/trauma-induced neuroimmune alterations and behavior. It is also worth noting that the lack of difference between remitted and current PTSD (in male-specific elevation of monocyte proportions) may have a number of implications for PTSD pathophysiology, including adverse health consequences associated with PTSD across the life course in men, which may be distinct from PTSD-associated health trajectories in women^118–120^.

Although the current dataset combined two cohorts and known pregnancies were excluded from our study, sample size and unavailable phenotype data on pregnancy, timing of the menstrual cycle, hormonal birth control use, and health status, as well as gender-related variables, such as coping mechanisms, are all limitations of this study. A small study in adult females that matched PTSD participants and controls for phase of menstrual cycle agreed with our female-specific results and reported no difference between PTSD subjects and controls in percentage of any lymphocyte subsets or total numbers of leukocytes, neutrophils, lymphocytes, or monocytes^121^. However, this study was small (PTSD: n = 16; controls: n= 15) and healthy controls did not have a history of trauma exposure. Future studies that account for hormone levels and other fundamental physiological sex differences may help identify female-specific associations between PTSD and leukocyte composition and clarify if hormone-dependent processes influence leukocyte composition dynamics.

Overall, our literature-supported finding of male-specific elevation in monocyte proportions illustrates feasibility of using DNAm-based leukocyte composition estimates to probe immune profiles. DNAm-based monocyte proportions may be an informative metric to include as part of a diagnostic biomarker panel for PTSD in males, and future study in females, with consideration for hormonal status, may elucidate a female-specific panel as well. Differential methylation markers discovered in sex-stratified EWAS, which account for these cell estimates as covariates, are other prime candidates to be included in such sex-specific biomarker panels. In fact, validation of recently published methods for determining cell-specific differential methylation profiles^122^ may enable the next significant advance in extracting insights from methylomic profiles by contextualizing how differential methylation in specific leukocyte subtypes alter regulatory dynamics in the immune system. Ultimately, this work may help to shape future studies designed to determine whether sex-specific methylomic metrics of peripheral immune status can inform us about sex differences in neuroinflammation and corresponding behavior in response to trauma exposure.

## 5. CONCLUSION

By combining DNA methylation datasets from two civilian cohorts, the current study found a small but significant elevation in monocyte proportion associated with lifetime PTSD in males, but not females. This sex-specific shift in peripheral blood leukocyte composition reflects a long-standing trait of PTSD diagnosis, rather than current state of PTSD. While this finding was confirmed using two different cell estimation approaches (i.e., deconvolution algorithms), the recently developed non-constrained projection approach (RPC) appears better suited for modeling leukocyte composition. Methylome-based characterization of immune profiles holds special promise for the study of PTSD and continued development of reference databases and validation of methods will build on these recent improvements to enrich our understanding of sex-specific immune dysregulation associated with PTSD.

## ACKNOWLEDGMENTS

We appreciate the time and effort of study participants, staff and volunteers of the Detroit Neighborhood Health Study, supported by the National Institutes of Health [R01DA022720, R01DA022720-S1, RC1MH088283, R01 MD011728, 3R01MD011728-02S1], and the Grady Trauma Project, supported by the National Institute of Mental Health [MH096764 and MH071537]. This work was also supported by the CompGen Fellowship (University of Illinois; GSK).

## LEGENDS FOR FIGURES AND TABLES

**Supplementary Figure 1:** Distribution of leukocyte subtypes based on constrained projection (CP) estimates, by sex. As noted in RPC estimates, sex differences in CD8^+^ T and CD56^+^ NK cell distributions were found to be prominent.

**Supplementary Figure 2:** Correlation (Spearman’s rho) between cell estimates derived using robust partial correlation (RPC) and constrained projection (CP) deconvolution algorithms is high in all leukocyte subtypes, with CD8^+^ T cells (CD8T) showing the worst agreement at *R* = 0.8 in females and *R* = 0.87 in males. CD8T cells also showed the largest discrepancy in sex-specific correlation of estimates. RPC-CP correlation was > 0.9 and difference in RPC-CP correlation between sexes was between 0.01 and 0.03 for all other leukocyte subtypes.

**Supplementary Figure 3:** Density plots for CP monocyte estimates in lifetime PTSD cases and controls, stratified by sex, show distinctly higher monocyte levels in males with lifetime PTSD compared to trauma-exposed cases. This difference in monocyte levels between lifetime PTSD case and controls is not observed in females, mirroring findings based on RPC estimates.

**Supplementary Figure 4:** Lifetime PTSD by sex interaction plot for estimated marginal means (EMMs) of CP monocyte estimates. Interaction plot shows a significant EMM difference between lifetime PTSD cases (red) and controls (blue) in males, where mean monocyte estimates are higher in cases than controls. No significant EMM difference was observed between PTSD cases and controls in females.

**Supplementary Figure 5:** Density plots for CP monocyte estimates comparing those with current PTSD, remitted PTSD, and trauma-exposed controls, stratified by sex. No significant difference is observed between current and remitted PTSD cases, which suggests that the significant peak shift in male PTSD cases is associated with long-standing PTSD trait, rather than current PTSD state. However, unlike RPC-based estimates, difference between PTSD case groups and trauma-exposed controls was only nominally significant in corresponding post-hoc Dunn test. No significant differences were observed in females.

**Supplementary Table 1:** Two-way ANCOVA Table for CP monocyte estimates (n = 469)

**Supplementary Table 2:** Summary for CP monocyte estimates by group This table describes untransformed CP monocyte estimates by group (i.e., sex and lifetime PTSD); n = count per group; EMM = estimated marginal means (i.e., least squares means); SE = standard errors for regular means; SE_EMM_ = standard errors for EMM. Lower and upper confidence limits are for 95% level. EMM and intervals were back-transformed from the square root scale to the original scale of cell subtype proportions (%). Significance level of alpha = 0.05 was used for EMM comparisons. Results for pairwise comparison were averaged over levels for current smoking. Degree of freedom was 461 and male lifetime PTSD cases were significantly different from other groups.

## REFERENCES

1. Association, A. P. Diagnostic and statistical manual of mental disorders. 5 edn, (2013).

2. Breslau, N. The epidemiology of trauma, PTSD, and other posttrauma disorders. Trauma Violence Abuse 10, 198–210, doi:10.1177/1524838009334448 (2009).

3. Kessler, R. C., Sonnega, A., Bromet, E., Hughes, M. & Nelson, C. B. Posttraumatic stress disorder in the National Comorbidity Survey. Arch Gen Psychiatry 52, 1048–1060 (1995).

4. Kilpatrick, D. G. et al. National estimates of exposure to traumatic events and PTSD prevalence using DSM-IV and DSM-5 criteria. J Trauma Stress 26, 537–547, doi:10.1002/jts.21848 (2013).

5. Liu, H. et al. Association of DSM-IV Posttraumatic Stress Disorder With Traumatic Experience Type and History in the World Health Organization World Mental Health Surveys. JAMA Psychiatry 74, 270–281, doi:10.1001/jamapsychiatry.2016.3783 (2017).

6. Benjet, C. et al. The epidemiology of traumatic event exposure worldwide: results from the World Mental Health Survey Consortium. Psychol Med 46, 327–343, doi:10.1017/S0033291715001981 (2016).

7. Kessler, R. C. Lifetime Prevalence and Age-of-Onset Distributions of DSM-IV Disorders in the National Comorbidity Survey Replication. Arch Gen Psychiatry 62, 593–602 (2005).

8. Kessler, R. C. & Wang, P. S. The descriptive epidemiology of commonly occurring mental disorders in the United States. Annu Rev Public Health 29, 115–129, doi:10.1146/annurev.publhealth.29.020907.090847 (2008).

9. Yehuda, R., Koenen, K. C., Galea, S. & Flory, J. D. The role of genes in defining a molecular biology of PTSD. Dis Markers 30, 67–76, doi:10.3233/DMA-2011-0794 (2011).

10. Breslau, N. et al. Trauma and Posttraumatic Stress Disorder in the Community. Arch Gen Psychiatry 55, 626–632. (1998).

11. Breslau, N., Davis, G. C., Andreski, P., Peterson, E. L. & Schultz, L. R. Sex differences in posttraumatic stress disorder. Arch Gen Psychiatry 54, 1044–1048 (1997).

12. Tolin, D. F. & Foa, E. B. Sex differences in trauma and posttraumatic stress disorder: a quantitative review of 25 years of research. Psychol Bull 132, 959–992, doi:10.1037/0033-2909.132.6.959 (2006).

13. Kessler, R. C., McGonagle, K. A., Swartz, M., Blazer, D. G. & Nelson, C. B. Sex and depression in the National Comorbidity Survey. I: Lifetime prevalence, chronicity and recurrence. J Affect Disord 29, 85–96 (1993).

14. Kessler, R. C. et al. Sex and depression in the National Comorbidity Survey. II: Cohort effects. J Affect Disord 30, 15–26 (1994).

15. Hodes, G. E. Sex, stress, and epigenetics: regulation of behavior in animal models of mood disorders. Biol Sex Differ 4, 1, doi:10.1186/2042-6410-4-1 (2013).

16. Bangasser, D. A. & Valentino, R. J. Sex differences in stress-related psychiatric disorders: neurobiological perspectives. Front Neuroendocrinol 35, 303–319, doi:10.1016/j.yfrne.2014.03.008 (2014).

17. Pooley, A. E. et al. Sex differences in the traumatic stress response: PTSD symptoms in women recapitulated in female rats. Biol Sex Differ 9, 31, doi:10.1186/s13293-018-0191-9 (2018).

18. Passos, I. C. et al. Inflammatory markers in post-traumatic stress disorder: a systematic review, meta-analysis, and meta-regression. The Lancet Psychiatry 2, 1002–1012, doi:10.1016/s2215-0366(15)00309-0 (2015).

19. Michopoulos, V., Powers, A., Gillespie, C. F., Ressler, K. J. & Jovanovic, T. Inflammation in Fear- and Anxiety-Based Disorders: PTSD, GAD, and Beyond. Neuropsychopharmacology 42, 254–270, doi:10.1038/npp.2016.146 (2017).

20. Wang, Z. & Young, M. R. PTSD, a Disorder with an Immunological Component. Front Immunol 7, 219, doi:10.3389/fimmu.2016.00219 (2016).

21. Kawamura, N., Kim, Y. & Asukai, N. Suppression of cellular immunity in men with a past history of posttraumatic stress disorder. Am J Psychiatry 158, 484–486, doi:10.1176/appi.ajp.158.3.484 (2001).

22. Altemus, M., Cloitre, M. & Dhabhar, F. S. Enhanced Cellular Immune Response in Women With PTSD Related to Childhood Abuse. Am J Psychiatry 160, 1705–1707 (2003).

23. Glover, D. A., Steele, A. C., Stuber, M. L. & Fahey, J. L. Preliminary evidence for lymphocyte distribution differences at rest and after acute psychological stress in PTSD-symptomatic women. Brain Behav Immun 19, 243–251, doi:10.1016/j.bbi.2004.08.002 (2005).

24. Gill, J. M., Saligan, L., Woods, S. & Page, G. PTSD is Associated With an Excess of Inflammatory Immune Activities. Perspectives in Psychiatric Care 45, 262–277 (2009).

25. Gotovac, K. et al. Natural killer cell cytotoxicity and lymphocyte perforin expression in veterans with posttraumatic stress disorder. Prog Neuropsychopharmacol Biol Psychiatry 34, 597–604, doi:10.1016/j.pnpbp.2010.02.018 (2010).

26. Lindqvist, D. et al. Proinflammatory milieu in combat-related PTSD is independent of depression and early life stress. Brain Behav Immun 42, 81–88, doi:10.1016/j.bbi.2014.06.003 (2014).

27. Aiello, A. E. et al. PTSD is associated with an increase in aged T cell phenotypes in adults living in Detroit. Psychoneuroendocrinology 67, 133–141, doi:10.1016/j.psyneuen.2016.01.024 (2016).

28. Lindqvist, D. et al. Increased pro-inflammatory milieu in combat related PTSD - A new cohort replication study. Brain Behav Immun 59, 260–264, doi:10.1016/j.bbi.2016.09.012 (2017).

29. Sondergaard, H. P., Hansson, L. O. & Theorell, T. The inflammatory markers C-reactive protein and serum amyloid A in refugees with and without posttraumatic stress disorder. Clin Chim Acta 342, 93–98, doi:10.1016/j.cccn.2003.12.019 (2004).

30. Sommershof, A. et al. Substantial reduction of naive and regulatory T cells following traumatic stress. Brain Behav Immun 23, 1117–1124, doi:10.1016/j.bbi.2009.07.003 (2009).

31. Hoge, E. A. et al. Broad spectrum of cytokine abnormalities in panic disorder and posttraumatic stress disorder. Depress Anxiety 26, 447–455, doi:10.1002/da.20564 (2009).

32. Michopoulos, V., Norrholm, S. D. & Jovanovic, T. Diagnostic Biomarkers for Posttraumatic Stress Disorder: Promising Horizons from Translational Neuroscience Research. Biol Psychiatry 78, 344–353, doi:10.1016/j.biopsych.2015.01.005 (2015).

33. Bersani, F. S. et al. A population of atypical CD56(-)CD16(+) natural killer cells is expanded in PTSD and is associated with symptom severity. Brain Behav Immun 56, 264–270, doi:10.1016/j.bbi.2016.03.021 (2016).

34. Wang, Z., Mandel, H., Levingston, C. A. & Young, M. R. I. An exploratory approach demonstrating immune skewing and a loss of coordination among cytokines in plasma and saliva of Veterans with combat-related PTSD. Hum Immunol 77, 652–657, doi:10.1016/j.humimm.2016.05.018 (2016).

35. Wohleb, E. S. et al. beta-Adrenergic receptor antagonism prevents anxiety-like behavior and microglial reactivity induced by repeated social defeat. J Neurosci 31, 6277–6288, doi:10.1523/JNEUROSCI.0450-11.2011 (2011).

36. Wohleb, E. S., Powell, N. D., Godbout, J. P. & Sheridan, J. F. Stress-induced recruitment of bone marrow-derived monocytes to the brain promotes anxiety-like behavior. J Neurosci 33, 13820–13833, doi:10.1523/JNEUROSCI.1671-13.2013 (2013).

37. Wohleb, E. S. et al. Re-establishment of anxiety in stress-sensitized mice is caused by monocyte trafficking from the spleen to the brain. Biol Psychiatry 75, 970–981, doi:10.1016/j.biopsych.2013.11.029 (2014).

38. McKim, D. B. et al. Sympathetic Release of Splenic Monocytes Promotes Recurring Anxiety Following Repeated Social Defeat. Biol Psychiatry 79, 803–813, doi:10.1016/j.biopsych.2015.07.010 (2016).

39. McKim, D. B. et al. Neuroinflammatory Dynamics Underlie Memory Impairments after Repeated Social Defeat. J Neurosci 36, 2590–2604, doi:10.1523/JNEUROSCI.2394-15.2016 (2016).

40. Wohleb, E. S. & Delpech, J. C. Dynamic cross-talk between microglia and peripheral monocytes underlies stress-induced neuroinflammation and behavioral consequences. Prog Neuropsychopharmacol Biol Psychiatry 79, 40–48, doi:10.1016/j.pnpbp.2016.04.013 (2017).

41. Bam, M. et al. Evidence for Epigenetic Regulation of Pro-Inflammatory Cytokines, Interleukin-12 and Interferon Gamma, in Peripheral Blood Mononuclear Cells from PTSD Patients. J Neuroimmune Pharmacol 11, 168–181, doi:10.1007/s11481-015-9643-8 (2016).

42. Zhou, J. et al. Dysregulation in microRNA expression is associated with alterations in immune functions in combat veterans with post-traumatic stress disorder. PLoS One 9, e94075, doi:10.1371/journal.pone.0094075 (2014).

43. Uddin, M. et al. Epigenetic and immune function profiles associated with posttraumatic stress disorder. Proc Natl Acad Sci U S A 107, 9470–9475, doi:10.1073/pnas.0910794107 (2010).

44. Rusiecki, J. A. et al. PTSD and DNA Methylation in Select Immune Function Gene Promoter Regions: A Repeated Measures Case-Control Study of U.S. Military Service Members. Front Psychiatry 4, 56, doi:10.3389/fpsyt.2013.00056 (2013).

45. Bam, M. et al. Dysregulated immune system networks in war veterans with PTSD is an outcome of altered miRNA expression and DNA methylation. Sci Rep 6, 31209, doi:10.1038/srep31209 (2016).

46. Smith, A. K. et al. Differential immune system DNA methylation and cytokine regulation in post-traumatic stress disorder. Am J Med Genet B Neuropsychiatr Genet 156B, 700–708, doi:10.1002/ajmg.b.31212 (2011).

47. Weaver, I. C. et al. Epigenetic programming by maternal behavior. Nat Neurosci 7, 847–854, doi:10.1038/nn1276 (2004).

48. McGowan, P. O. et al. Epigenetic regulation of the glucocorticoid receptor in human brain associates with childhood abuse. Nat Neurosci 12, 342–348, doi:10.1038/nn.2270 (2009).

49. Stankiewicz, A. M., Swiergiel, A. H. & Lisowski, P. Epigenetics of stress adaptations in the brain. Brain Res Bull 98, 76–92, doi:10.1016/j.brainresbull.2013.07.003 (2013).

50. Miller, C. A., Campbell, S. L. & Sweatt, J. D. DNA methylation and histone acetylation work in concert to regulate memory formation and synaptic plasticity. Neurobiol Learn Mem 89, 599–603, doi:10.1016/j.nlm.2007.07.016 (2008).

51. Maddox, S. A., Schafe, G. E. & Ressler, K. J. Exploring epigenetic regulation of fear memory and biomarkers associated with post-traumatic stress disorder. Front Psychiatry 4, 62, doi:10.3389/fpsyt.2013.00062 (2013).

52. Zovkic, I. B. & Sweatt, J. D. Epigenetic mechanisms in learned fear: implications for PTSD. Neuropsychopharmacology 38, 77–93, doi:10.1038/npp.2012.79 (2013).

53. Malan-Muller, S., Seedat, S. & Hemmings, S. M. Understanding posttraumatic stress disorder: insights from the methylome. Genes Brain Behav 13, 52–68, doi:10.1111/gbb.12102 (2014).

54. Kwapis, J. L. & Wood, M. A. Epigenetic mechanisms in fear conditioning: implications for treating post-traumatic stress disorder. Trends Neurosci 37, 706–720, doi:10.1016/j.tins.2014.08.005 (2014).

55. Alvarez-Errico, D., Vento-Tormo, R., Sieweke, M. & Ballestar, E. Epigenetic control of myeloid cell differentiation, identity and function. Nat Rev Immunol 15, 7–17, doi:10.1038/nri3777 (2015).

56. Sun, B. et al. DNA methylation perspectives in the pathogenesis of autoimmune diseases. Clin Immunol 164, 21–27, doi:10.1016/j.clim.2016.01.011 (2016).

57. Chen, L. et al. Genetic Drivers of Epigenetic and Transcriptional Variation in Human Immune Cells. Cell 167, 1398–1414 e1324, doi:10.1016/j.cell.2016.10.026 (2016).

58. Wolf, E. J. et al. Traumatic stress and accelerated DNA methylation age: A meta-analysis. Psychoneuroendocrinology, doi:10.1016/j.psyneuen.2017.12.007 (2017).

59. Irwin, M. R. & Cole, S. W. Reciprocal regulation of the neural and innate immune systems. Nat Rev Immunol 11, 625–632, doi:10.1038/nri3042 (2011).

60. Pfau, M. L. & Russo, S. J. Peripheral and Central Mechanisms of Stress Resilience. Neurobiol Stress 1, 66–79, doi:10.1016/j.ynstr.2014.09.004 (2015).

61. Titus, A. J., Gallimore, R. M., Salas, L. A. & Christensen, B. C. Cell-type deconvolution from DNA methylation: a review of recent applications. Hum Mol Genet 26, R216–R224, doi:10.1093/hmg/ddx275 (2017).

62. Teschendorff, A. E., Breeze, C. E., Zheng, S. C. & Beck, S. A comparison of reference-based algorithms for correcting cell-type heterogeneity in Epigenome-Wide Association Studies. BMC Bioinformatics 18, 105, doi:10.1186/s12859-017-1511-5 (2017).

63. Koestler, D. C. et al. Improving cell mixture deconvolution by identifying optimal DNA methylation libraries (IDOL). BMC Bioinformatics 17, 120, doi:10.1186/s12859-016-0943-7 (2016).

64. Salas, L. A. et al. An optimized library for reference-based deconvolution of whole-blood biospecimens assayed using the Illumina HumanMethylationEPIC BeadArray. Genome Biol 19, 64, doi:10.1186/s13059-018-1448-7 (2018).

65. Wiencke, J. K. et al. Immunomethylomic approach to explore the blood neutrophil lymphocyte ratio (NLR) in glioma survival. Clin Epigenetics 9, 10, doi:10.1186/s13148-017-0316-8 (2017).

66. Koestler, D. C. et al. DNA Methylation-Derived Neutrophil-to-Lymphocyte Ratio: An Epigenetic Tool to Explore Cancer Inflammation and Outcomes. Cancer Epidemiol Biomarkers Prev 26, 328–338, doi:10.1158/1055-9965.EPI-16-0461 (2017).

67. Goldmann, E. et al. Pervasive exposure to violence and posttraumatic stress disorder in a predominantly African American Urban Community: the Detroit Neighborhood Health Study. J Trauma Stress 24, 747–751, doi:10.1002/jts.20705 (2011).

68. Meyers, J. L. et al. Frequency of alcohol consumption in humans; the role of metabotropic glutamate receptors and downstream signaling pathways. Transl Psychiatry 5, e586, doi:10.1038/tp.2015.70 (2015).

69. Gillespie, C. F. et al. Trauma exposure and stress-related disorders in inner city primary care patients. Gen Hosp Psychiatry 31, 505–514, doi:10.1016/j.genhosppsych.2009.05.003 (2009).

70. Binder, E. Association of FKBP5 Polymorphisms and Childhood Abuse With Risk of Posttraumatic Stress Disorder Symptoms in Adults. JAMA 299 (2008).

71. Zannas, A. S. et al. Lifetime stress accelerates epigenetic aging in an urban, African American cohort: relevance of glucocorticoid signaling. Genome Biol 16, 266, doi:10.1186/s13059-015-0828-5 (2015).

72. Luppi, P. How immune mechanisms are affected by pregnancy. Vaccine 21, 3352–3357, doi:10.1016/S0264-410x(03)00331-1 (2003).

73. Blanchard, E. B. Psychometric properties of the PTSD checklist (PCL). Behav Res Ther 34, 669–673 (1996).

74. Grubaugh, A. L., Elhai, J. D., Cusack, K. J., Wells, C. & Frueh, B. C. Screening for PTSD in public-sector mental health settings: the diagnostic utility of the PTSD checklist. Depress Anxiety 24, 124–129, doi:10.1002/da.20226 (2007).

75. Wilkins, K. C., Lang, A. J. & Norman, S. B. Synthesis of the psychometric properties of the PTSD checklist (PCL) military, civilian, and specific versions. Depress Anxiety 28, 596–606, doi:10.1002/da.20837 (2011).

76. Ruggiero, K. J., Del Ben, K., Scotti, J. R. & Rabalais, A. E. Psychometric properties of the PTSD Checklist-Civilian Version. J Trauma Stress 16, 495–502, doi:10.1023/A:1025714729117 (2003).

77. Koenen, K. C. et al. SLC6A4 methylation modifies the effect of the number of traumatic events on risk for posttraumatic stress disorder. Depress Anxiety 28, 639–647, doi:10.1002/da.20825 (2011).

78. Uddin, M. et al. Gene expression and methylation signatures of MAN2C1 are associated with PTSD. Dis Markers 30, 111–121, doi:10.3233/DMA-2011-0750 (2011).

79. Blake, D. D. et al. The development of a Clinician-Administered PTSD Scale. J Trauma Stress 8, 75–90 (1995).

80. Ratanatharathorn, A. et al. Epigenome-wide association of PTSD from heterogeneous cohorts with a common multi-site analysis pipeline. Am J Med Genet B Neuropsychiatr Genet 174, 619–630, doi:10.1002/ajmg.b.32568 (2017).

81. Uddin, M. et al. Epigenetic meta-analysis across three civilian cohorts identifies NRG1 and HGS as blood-based biomarkers for post-traumatic stress disorder. Epigenomics, doi:10.2217/epi-2018-0049 (2018).

82. Mehta, D. et al. Childhood maltreatment is associated with distinct genomic and epigenetic profiles in posttraumatic stress disorder. Proc Natl Acad Sci U S A 110, 8302–8307, doi:10.1073/pnas.1217750110 (2013).

83. R: A Language and Environment for Statistical Computing (R Foundation for Statistical Computing, Vienna, Austria, 2018).

84. Aryee, M. J. et al. Minfi: a flexible and comprehensive Bioconductor package for the analysis of Infinium DNA methylation microarrays. Bioinformatics 30, 1363–1369, doi:10.1093/bioinformatics/btu049 (2014).

85. Huber, W. et al. Orchestrating high-throughput genomic analysis with Bioconductor. Nat Methods 12, 115–121, doi:10.1038/nmeth.3252 (2015).

86. Gentleman, R. C. et al. Bioconductor: open software development for computational biology and bioinformatics. Genome Biol 5, R80, doi:10.1186/gb-2004-5-10-r80 (2004).

87. Barfield, R. T., Kilaru, V., Smith, A. K. & Conneely, K. N. CpGassoc: an R function for analysis of DNA methylation microarray data. Bioinformatics 28, 1280–1281, doi:10.1093/bioinformatics/bts124 (2012).

88. Xu, Z., Niu, L., Li, L. & Taylor, J. A. ENmix: a novel background correction method for Illumina HumanMethylation450 BeadChip. Nucleic Acids Res 44, e20, doi:10.1093/nar/gkv907 (2016).

89. Liu, J. & Siegmund, K. D. An evaluation of processing methods for HumanMethylation450 BeadChip data. BMC Genomics 17, 469, doi:10.1186/s12864-016-2819-7 (2016).

90. Fortin, J. P., Triche, T. J., Jr. & Hansen, K. D. Preprocessing, normalization and integration of the Illumina HumanMethylationEPIC array with minfi. Bioinformatics 33, 558–560, doi:10.1093/bioinformatics/btw691 (2017).

91. Chen, Y. A. et al. Discovery of cross-reactive probes and polymorphic CpGs in the Illumina Infinium HumanMethylation450 microarray. Epigenetics 8, 203–209, doi:10.4161/epi.23470 (2013).

92. Teschendorff, A. E. et al. A beta-mixture quantile normalization method for correcting probe design bias in Illumina Infinium 450 k DNA methylation data. Bioinformatics 29, 189–196, doi:10.1093/bioinformatics/bts680 (2013).

93. Morris, T. J. et al. ChAMP: 450k Chip Analysis Methylation Pipeline. Bioinformatics 30, 428–430, doi:10.1093/bioinformatics/btt684 (2014).

94. Tian, Y. et al. ChAMP: updated methylation analysis pipeline for Illumina BeadChips. Bioinformatics 33, 3982–3984, doi:10.1093/bioinformatics/btx513 (2017).

95. Triche, T. J., Jr, Weisenberger, D. J., Van Den Berg, D., Laird, P. W. & Siegmund, K. D. Low-level processing of Illumina Infinium DNA Methylation BeadArrays. Nucleic Acids Res 41, e90, doi:10.1093/nar/gkt090 (2013).

96. Johnson, W. E., Li, C. & Rabinovic, A. Adjusting batch effects in microarray expression data using empirical Bayes methods. Biostatistics 8, 118–127, doi:10.1093/biostatistics/kxj037 (2007).

97. Leek, J. T. The SVA package for removing batch effects and other unwanted variation in high-throughput experiments. (2012).

98. Roadmap Epigenomics, C. et al. Integrative analysis of 111 reference human epigenomes. Nature 518, 317–330, doi:10.1038/nature14248 (2015).

99. Houseman, E. A. DNA methylation arrays as surrogate measures of cell mixture distribution. BMC Bioinformatics 13 (2012).

100. Rahmani, E. et al. Genome-wide methylation data mirror ancestry information. Epigenetics Chromatin 10, 1, doi:10.1186/s13072-016-0108-y (2017).

101. Barfield, R. T. et al. Accounting for population stratification in DNA methylation studies. Genet Epidemiol 38, 231–241, doi:10.1002/gepi.21789 (2014).

102. Scharer, C. D., Barwick, B. G., Youngblood, B. A., Ahmed, R. & Boss, J. M. Global DNA methylation remodeling accompanies CD8 T cell effector function. J Immunol 191, 3419–3429, doi:10.4049/jimmunol.1301395 (2013).

103. Inoshita, M. et al. Sex differences of leukocytes DNA methylation adjusted for estimated cellular proportions. Biol Sex Differ 6, 11, doi:10.1186/s13293-015-0029-7 (2015).

104. Abdullah, M. et al. Gender effect on in vitro lymphocyte subset levels of healthy individuals. Cell Immunol 272, 214–219, doi:10.1016/j.cellimm.2011.10.009 (2012).

105. White, N. et al. Accounting for cell lineage and sex effects in the identification of cell-specific DNA methylation using a Bayesian model selection algorithm. PLoS One 12, e0182455, doi:10.1371/journal.pone.0182455 (2017).

106. Suarez-Alvarez, B., Rodriguez, R. M., Fraga, M. F. & Lopez-Larrea, C. DNA methylation: a promising landscape for immune system-related diseases. Trends Genet 28, 506–514, doi:10.1016/j.tig.2012.06.005 (2012).

107. Johnson, G. J., Slater, B. C., Leis, L. A., Rector, T. S. & Bach, R. R. Blood Biomarkers of Chronic Inflammation in Gulf War Illness. PLoS One 11, e0157855, doi:10.1371/journal.pone.0157855 (2016).

108. Engler, H., Bailey, M. T., Engler, A. & Sheridan, J. F. Effects of repeated social stress on leukocyte distribution in bone marrow, peripheral blood and spleen. J Neuroimmunol 148, 106–115, doi:10.1016/j.jneuroim.2003.11.011 (2004).

109. Weber, M. D., Godbout, J. P. & Sheridan, J. F. Repeated Social Defeat, Neuroinflammation, and Behavior: Monocytes Carry the Signal. Neuropsychopharmacology 42, 46–61, doi:10.1038/npp.2016.102 (2017).

110. Niraula, A., Witcher, K. G., Sheridan, J. F. & Godbout, J. P. Interleukin-6 Induced by Social Stress Promotes a Unique Transcriptional Signature in the Monocytes That Facilitate Anxiety. Biological Psychiatry, doi:10.1016/j.biopsych.2018.09.030 (2018).

111. Bekhbat, M. & Neigh, G. N. Sex differences in the neuro-immune consequences of stress: Focus on depression and anxiety. Brain Behav Immun 67, 1–12, doi:10.1016/j.bbi.2017.02.006 (2018).

112. Bekhbat, M. & Neigh, G. N. Stress-induced neuroimmune priming in males and females: Comparable but not identical. Brain Behav Immun 73, 149–150, doi:10.1016/j.bbi.2018.05.001 (2018).

113. O’Donovan, A. et al. Transcriptional control of monocyte gene expression in post-traumatic stress disorder. Dis Markers 30, 123–132, doi:10.3233/DMA-2011-0768 (2011).

114. Neylan, T. C. et al. Suppressed monocyte gene expression profile in men versus women with PTSD. Brain Behav Immun 25, 524–531, doi:10.1016/j.bbi.2010.12.001 (2011).

115. Nunn, C. L., Lindenfors, P., Pursall, E. R. & Rolff, J. On sexual dimorphism in immune function. Philos Trans R Soc Lond B Biol Sci 364, 61–69, doi:10.1098/rstb.2008.0148 (2009).

116. Klein, S. L. & Flanagan, K. L. Sex differences in immune responses. Nat Rev Immunol 16, 626–638, doi:10.1038/nri.2016.90 (2016).

117. Scotland, R. S., Stables, M. J., Madalli, S., Watson, P. & Gilroy, D. W. Sex differences in resident immune cell phenotype underlie more efficient acute inflammatory responses in female mice. Blood 118, 5918–5927, doi:10.1182/blood-2011-03-340281 (2011).

118. Dedert, E. A., Calhoun, P. S., Watkins, L. L., Sherwood, A. & Beckham, J. C. Posttraumatic stress disorder, cardiovascular, and metabolic disease: a review of the evidence. Ann Behav Med 39, 61–78, doi:10.1007/s12160-010-9165-9 (2010).

119. Mitchell, K. S. et al. PTSD and obesity in the Detroit neighborhood health study. Gen Hosp Psychiatry 35, 671–673, doi:10.1016/j.genhosppsych.2013.07.015 (2013).

120. McLean, C. P., Asnaani, A., Litz, B. T. & Hofmann, S. G. Gender differences in anxiety disorders: prevalence, course of illness, comorbidity and burden of illness. J Psychiatr Res 45, 1027–1035, doi:10.1016/j.jpsychires.2011.03.006 (2011).

121. Altemus, M., Dhabhar, F. S. & Yang, R. Immune function in PTSD. Ann N Y Acad Sci 1071, 167–183, doi:10.1196/annals.1364.013 (2006).

122. Zheng, S. C., Breeze, C. E., Beck, S. & Teschendorff, A. E. Identification of differentially methylated cell types in epigenome-wide association studies. Nat Methods 15, 1059–1066, doi:10.1038/s41592-018-0213-x (2018).

