## Supplemental Material for "Methylomic profiles reveal sex-specific shifts in leukocyte composition associated with post-traumatic stress disorder"

**Supplementary Figure 1:** Distribution of leukocyte subtypes based on constrained projection (CP) estimates, by sex. As noted in RPC estimates, sex differences in $CD8^{+}$ T and $CD56^{+}$ NK cell distributions were found to be prominent.

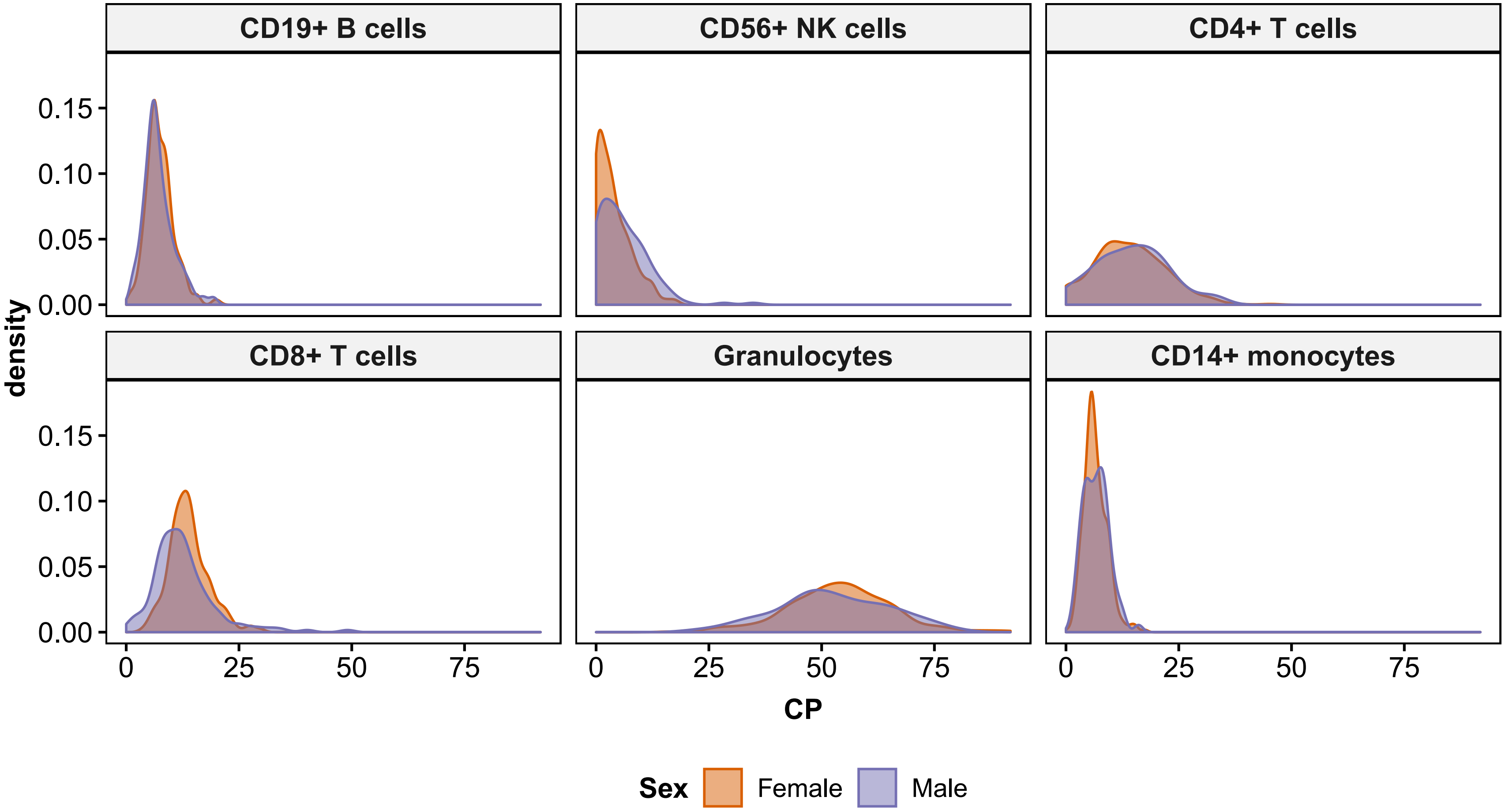

**Supplementary Figure 2:** Correlation (Spearman's rho) between cell estimates derived using robust partial correlation (RPC) and constrained projection (CP) deconvolution algorithms is high in all leukocyte subtypes, with CD8$^{+}$ T cells (CD8T) showing the worst agreement at $R = 0.8$ in females and $R = 0.87$ in males. CD8T cells also showed the largest discrepancy in sex-specific correlation of estimates. RPC-CP correlation was > 0.9 and difference in RPC-CP correlation between sexes was between 0.01 and 0.03 for all other leukocyte subtypes.

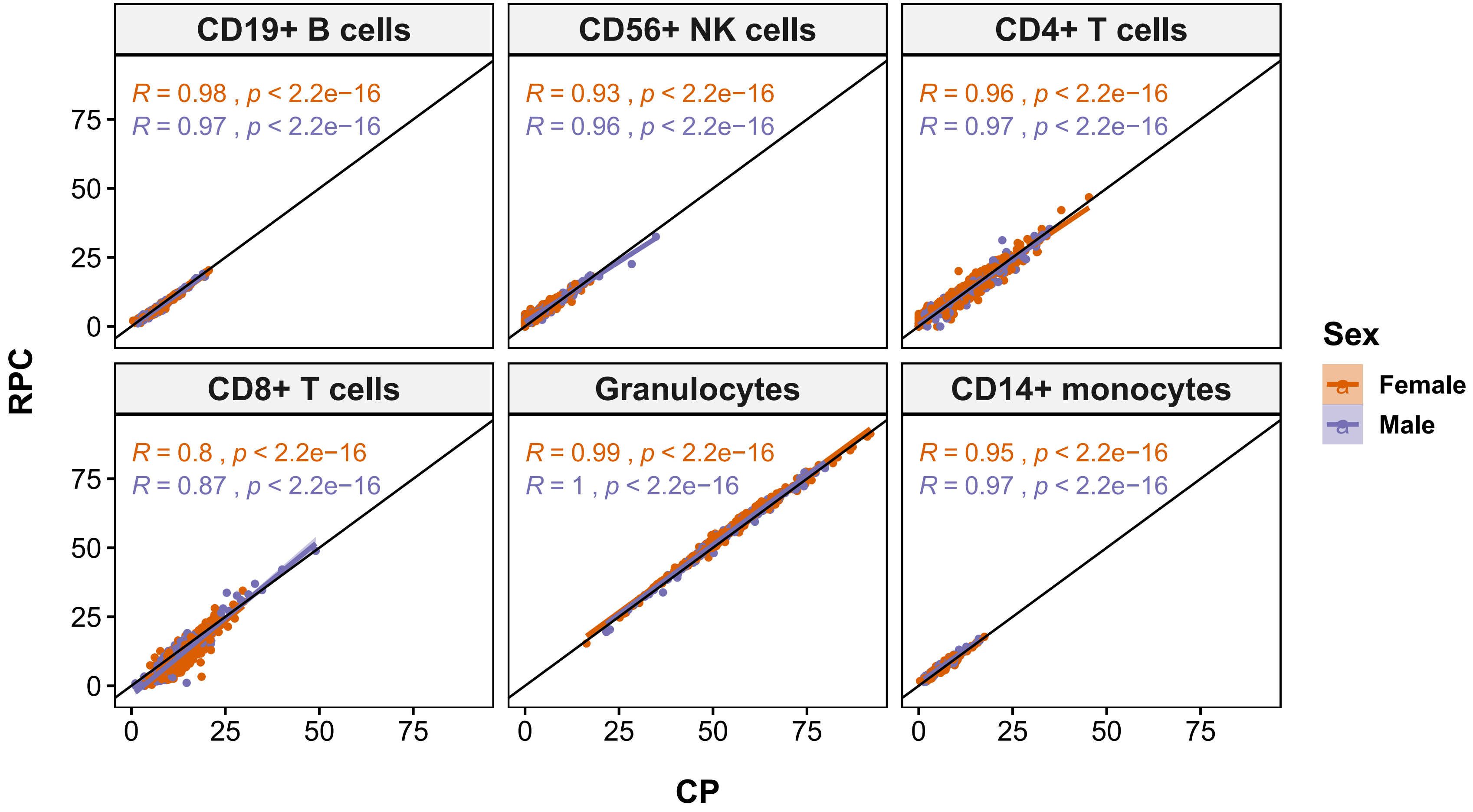

**Supplementary Figure 3:** Density plots for CP monocyte estimates in lifetime PTSD cases and controls, stratified by sex, show distinctly higher monocyte levels in males with lifetime PTSD compared to trauma-exposed cases.  This difference in monocyte levels between lifetime PTSD case and controls is not observed in females, mirroring findings based on RPC estimates.

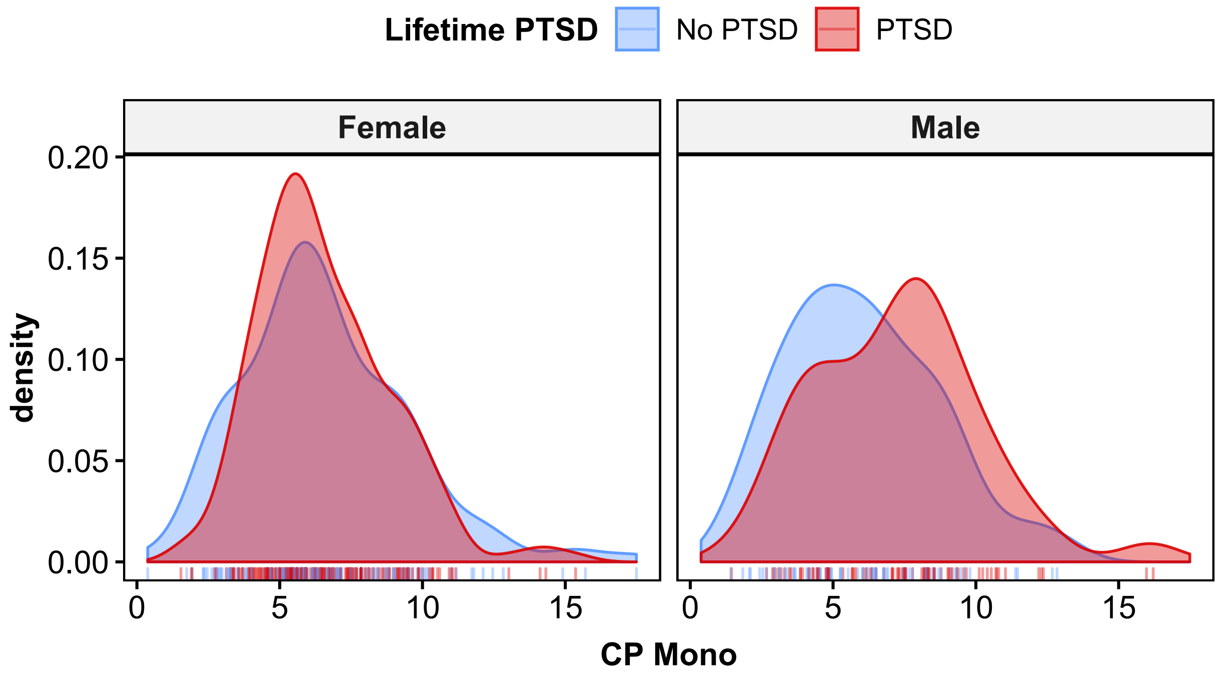

**Supplementary Figure 4:** Lifetime PTSD by sex interaction plot for estimated marginal means (EMMs) of CP monocyte estimates. Interaction plot shows a significant EMM difference between lifetime PTSD cases (red) and controls (blue) in males, where mean monocyte estimates are higher in cases than controls. No significant EMM difference was observed between PTSD cases and controls in females.

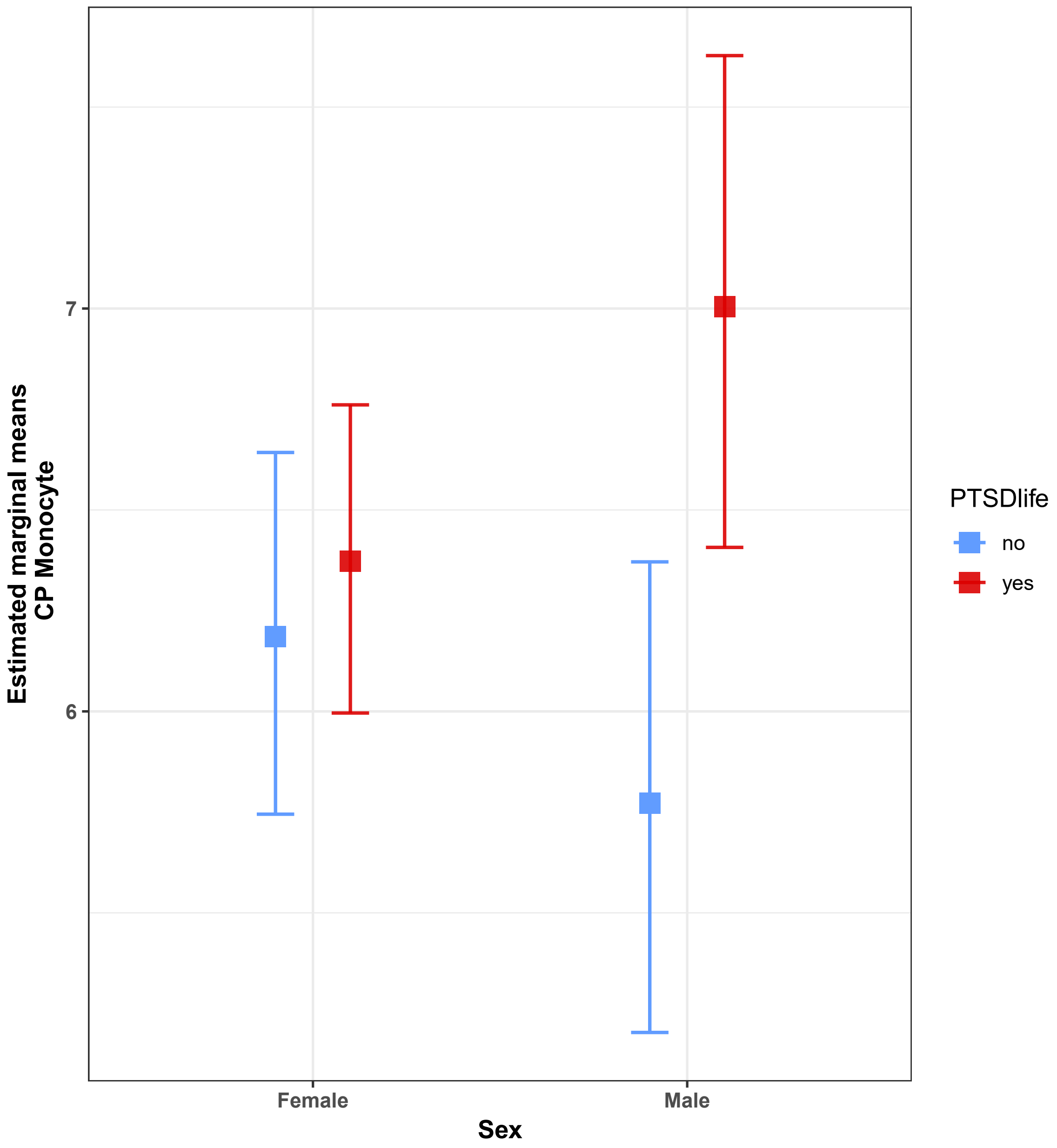

**Supplementary Figure 5:** Density plots for CP monocyte estimates comparing those with current PTSD, remitted PTSD, and trauma-exposed controls, stratified by sex. No significant difference is observed between current and remitted PTSD cases, which suggests that the significant peak shift in male PTSD cases is associated with long-standing PTSD trait, rather than current PTSD state. However, unlike RPC-based estimates, difference between PTSD case groups and trauma-exposed controls was only nominally significant in corresponding post-hoc Dunn test. No significant differences were observed in females.

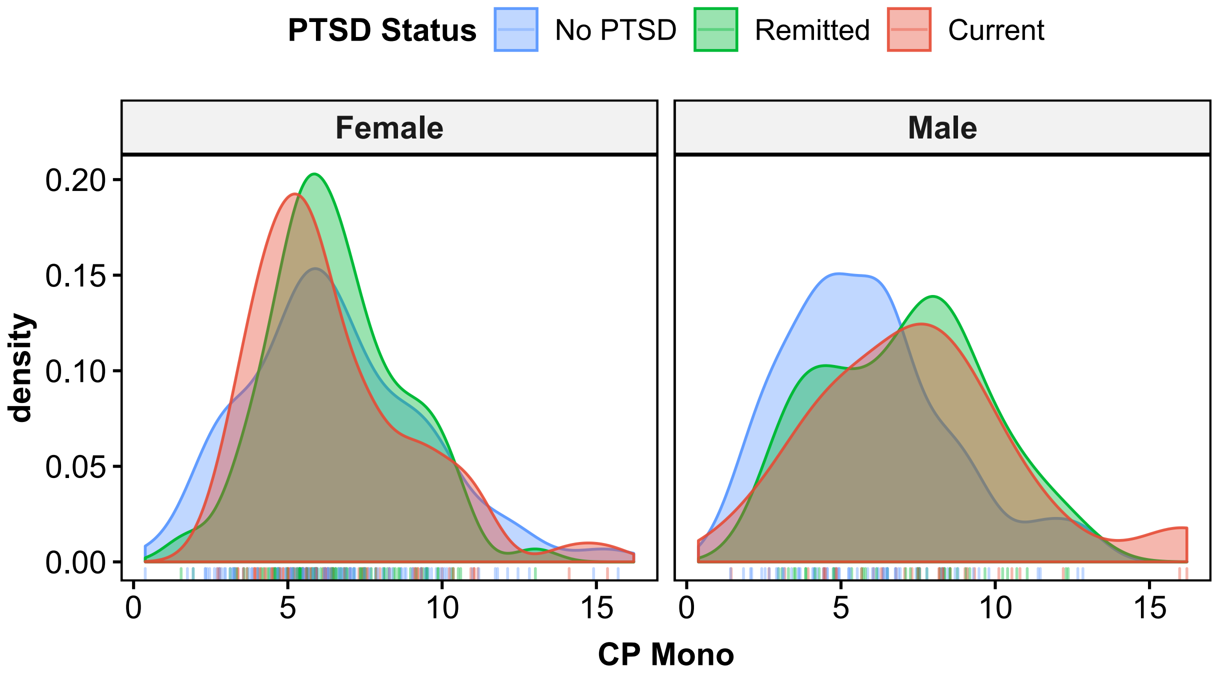

**Supplementary Table 1:** Two-way ANCOVA Table for CP monocyte estimates (n = 469)

| Terms | Type III Sum of Squares | *df* | Mean Square | *F* | *p* | partial $\eta^2$ |
|---|---|---|---|---|---|---|
| Sex | 0.034 | 1 | 0.034 | 0.130 | 0.719 | 0.000 |
| PTSDlife | 1.929 | 1 | 1.929 | 7.256 | 0.007** | 0.015 |
| Age | 2.964 | 1 | 2.964 | 11.150 | 0.001*** | 0.024 |
| ancPC1 | 0.036 | 1 | 0.036 | 0.137 | 0.712 | 0.000 |
| ancPC2 | 0.063 | 1 | 0.063 | 0.236 | 0.627 | 0.001 |
| Smoking | 0.000 | 1 | 0.000 | 0.000 | 0.987 | 0.000 |
| Sex:PTSDlife | 1.060 | 1 | 1.060 | 3.987 | 0.046* | 0.009 |
| Residuals | 122.555 | 461 | 0.266 | | | |

** $p < 0.05$, ** $p < 0.01$, *** $p < 0.005$

**Supplementary Table 2:** Summary for CP monocyte estimates by group

| Sex | PTSD | n | mean | SE | EMM | $SE_{EMM}$ | lower.CL | upper.CL |
|---|---|---|---|---|---|---|---|---|
| Female | no | 135 | 6.564 | 0.2499 | 6.186 | 0.2285 | 5.745 | 6.643 |
| Female | yes | 184 | 6.551 | 0.1735 | 6.373 | 0.1946 | 5.996 | 6.761 |
| Male | no | 70 | 6.072 | 0.3165 | 5.772 | 0.2972 | 5.203 | 6.371 |
| Male | yes | 80 | 7.270 | 0.3248 | 7.004 | 0.3106 | 6.407 | 7.628 |

This table describes untransformed CP monocyte estimates by group (i.e., sex and lifetime PTSD); n = count per group; EMM = estimated marginal means (i.e., least squares means); SE = standard errors for regular means; $SE_{EMM}$ = standard errors for EMM.

Lower and upper confidence limits are for 95% level. EMM and intervals were back-transformed from the square root scale to the original scale of cell subtype proportions (%). Significance level of alpha = 0.05 was used for EMM comparisons. Results for pairwise comparison were averaged over levels for current smoking. Degree of freedom was 461 and male lifetime PTSD cases were significantly different from other groups.
